# Spatiotemporal coordination of Rac1 and Cdc42 at the whole cell level during cell ruffling

**DOI:** 10.1101/2023.03.31.535147

**Authors:** Siarhei Hladyshau, Jorik P Stoop, Kosei Kamada, Shuyi Nie, Denis V Tsygankov

**Affiliations:** School of Biology, Georgia Institute of Technology, Atlanta, GA, USA; Wallace H. Coulter Department of Biomedical Engineering, Georgia Institute of Technology and Emory University, Atlanta, GA, USA; Faculty of Medicine, The University of Tokyo, Tokyo, Japan

**Keywords:** Multiscale modeling, morphodynamics, Rho family GTPases, FRET-based biosensors, cytoskeletal regulation

## Abstract

Rho-GTPases are central regulators within a complex signaling network that controls the cytoskeletal organization and cell movement. This network includes multiple GTPases, such as the most studied Rac1, Cdc42, and RhoA, and their numerous effectors that provide mutual regulation and feedback loops. Here we investigate the temporal and spatial relationship between Rac1 and Cdc42 during membrane ruffling using a simulation model which couples GTPase signaling with cell morphodynamics to capture the GTPase behavior observed with FRET-based biosensors. We show that membrane velocity is regulated by the kinetic rate of GTPase activation rather than the concentration of active GTPase. Our model captures both uniform and polarized ruffling. We also show that cell-type specific time delays between Rac1 and Cdc42 activation can be reproduced with a single signaling motif, in which the delay is controlled by feedback from Cdc42 to Rac1. The resolution of our simulation output matches those of the time-lapsed recordings of cell dynamics and GTPase activity. This approach allows us to validate simulation results with quantitative precision using the same pipeline for the analysis of simulated and experimental data.

## Introduction

Cell motion and change in shape (also referred to as morphodynamics) is a process driven by complex, multiscale machinery of the cytoskeleton. The central regulators of this machinery are small GTPases that control both membrane protrusion and retraction. GTPases control protrusion by activating F-actin polymerization through nucleation-promoting factors (downstream of GTPases Rac1 and Cdc42) [1], and they modulate retraction by controlling actomyosin contractility through Rho kinase (downstream of GTPase RhoA) [2]. Over the last decades, numerous studies have shed light on the mechanisms of RhoA, Rac1, and Cdc42 signaling (reviewed in [3-8]). Many regulators of GTPase signaling were discovered, including guanine nucleotide exchange factors (GEFs) that control GTPase activation by catalyzing the exchange of GDP with GTP, and guanine activating proteins (GAPs) that control GTPase deactivation by catalyzing the GTP hydrolysis [4]. Some of these proteins were shown to interact with several GTPases or have dual GEF/GAP functions [7, 9-11]. Such multi-target regulators may work as a crosslink between Rac1 and Cdc42. Indeed, it was reported that induced activation of Cdc42 can lead to the activation of Rac1, but the specific regulators and the details of this mechanism are not fully understood [12]. Similarly, the crosstalk of small GTPases with other signaling pathways is a matter of intensive research [13]. It was reported that signaling pathways related to the cytoskeleton regulation, including the Rac1 pathway, operate downstream of an excitable signal transduction network that involves Ras and PI3K pathways [14, 15]. Given that many effectors of small GTPases have a pleckstrin homology (PH) domain, which binds to phosphatidylinositol lipids, Rac1 and Ras/PI3K signaling networks can be coupled in multiple ways (e.g., through interactions of Rac1 GEFs with PIP3) [13]. Also, the crosstalk between phosphoinositides and Cdc42 was reported in [16]. Despite the growing knowledge of the roles of various cytoskeletal regulators, there is still no holistic understanding of the intertwined small-GTPase signaling pathways to this day. This lack of understanding is especially prevalent in the context of their coordinated activity during cell motion and interaction with the extracellular environment.

The application of fluorescence resonance energy transfer (FRET) biosensors has facilitated the study of small GTPases and their effectors in live cells with high spatial and temporal resolution [17]. This technology has allowed researchers to investigate the quantitative relationship between the proteins’ activity and the velocity of membrane protrusion [17, 18]. Several studies showed that during protrusion, the velocity peak precedes the peak of Rac1 and Cdc42 activity [19, 20]. This appears to be counterintuitive as the F-actin polymerization that drives protrusion is regulated downstream of these GTPases. However, Yamao *et al*. showed that the response function between biosensor activity and membrane velocity displays the properties of a differentiator circuit [21]. The authors of this study suggested that membrane dynamics is regulated by the temporal derivative of active forms of Rac1 and Cdc42, which may explain the time shift between membrane velocity and GTPase activity. Yet, the mechanism that would explain how cell membrane protrusion could be regulated by the temporal derivative of GTPase activity has not been described. Using cross-correlation analysis, Marston *et al*. showed that in the epithelial breast cancer cell line (MDA-MB-231), there is no measurable time shift between the peaks of Cdc42 and Rac1 activities [17]. On the other hand, some studies reported that Rac1 activation could be induced by Cdc42 [3, 12] and that timing between the activation of two GTPases may vary [22, 23]. These results suggest that the relative timing and regulation of GTPase may be cell-type specific and depend on the biological context or type of regulation. Partial correlation analysis also revealed that the GEF Asef contributes to the regulation of cell protrusion by both Cdc42 and Rac1, but the degree of the GEF’s influence on the two GTPase pathways is different [17]. This computational analysis takes advantage of the simultaneous visualization of two GTPases or a GTPase and its regulator and provides valuable insights regarding mutual influences between signaling components. However, correlation analysis doesn’t provide the full mechanistic picture of a regulatory process. Thus, in this work, we seek to develop a model that couples GTPase signaling and cell edge motion. We quantitatively analyze the relationships of GTPase activity level and its rate of change with cell edge position and velocity to directly compare the experimental data and our simulation results.

The two key features characteristic of models used in the theory of biological morphogenesis are autocatalytic activation of the pattern-forming component and significant difference in the diffusion coefficient of an activator and an inhibitor [24-27]. This theory was originally proposed by Alan Turing [28] and further developed by Hans Meinhardt [24, 29, 30]. Later, researchers applied these developments to model small GTPases with the assumption of an underlying activator-inhibitor mechanism that allows for representing their localized activation and wave dynamics [31, 32]. Indeed, GTPases exhibit the properties of autocatalytic activation through positive feedback [33-36] and a significant difference in the diffusion coefficients of active and inactive forms due to the interaction of active GTPases with the membrane [11, 37]. While activator-inhibitor models are consistent with the observed features of GTPase activity in cells, it is hard to relate the model parameters to the biochemical characteristics of the components in the signaling motif [14, 38]. Recent studies proposed more detailed models to improve the interpretability of the modeling results [9]. Specifically, to account for the switch of small GTPases between active and inactive states, the mass-conserved reaction-diffusion model (MCRD, also known as the wave-pinning mechanism) was proposed as a framework to study the quantitative properties of GTPase signaling [27, 39]. It was successfully applied to study cell polarization [27, 40-43]. Importantly, this model captures the Turing-unstable type of behavior, in which the activity of small-GTPases remains non-homogeneous, as well as the transition to the excitable regime, in which a stimulus-induced activation is required to generate GTPase patterning. Therefore, the MCRD model can be used to represent a core signaling motif of GTPases and, when coupled with downstream and upstream effectors, can reproduce rich dynamics, including the formation of complex cortical waves of GTPase activity [44]. A more detailed variation of this model was also successfully applied to study the coordination of RhoA and Rac1 in the leading and trailing edges of moving cells [45]. A similar modeling method was applied to analyze drug resistance to cancer treatment [46]. Thus, the MCRD-based modeling framework is emerging as a powerful quantitative approach to studying the mechanisms of coordinated spatiotemporal behavior of small-GTPases and their effects on cell morphodynamics.

In this work, we focus our modeling effort on investigating the regulation of Rac1 and Cdc42 in the epithelial breast cancer cell line (MDA-MB-231) during cell membrane ruffling. We used experimental data from a previously published study on Rac1 and Cdc42 regulation with multiplexed FRET biosensors [17]. In these experiments, cell morphodynamics typically involves two timescales: fast localized oscillations of the cell outline and a slow change of the overall cell shape. To account for these features of the experimental data and provide accurate measurements of the protrusive activity, we developed an automated image analysis pipeline. This pipeline tracks fast cell edge movement and local biosensor signals along many evenly distributed line segments, which slowly move together with the overall cell shape. To understand the regulatory mechanism of such dynamics, we developed a computational framework to model cell morphodynamics coupled with a reaction-diffusion representation of GTPase activity in the cell. Using this framework, we showed that regulation of cell edge velocity by kinetic rates of GTPase activation accurately reproduces the relationship between the protrusive activity and the biosensor signal in experimental data, consistent with the differentiator circuit interpretation. Using our image analysis pipeline, we also analyzed the mouse-embryonic fibroblasts (MEFs) data, published previously by MacNevin *et al*. [47], and observed a delayed activation of Rac1 relatively Cdc42. Although we obtained a delay of 5 seconds, which is at the limit of the temporal resolution of the available imaging data, we cannot exclude that such a delay is a true feature of the cell edge motion in MEFs. An experiment with a higher frame rate is needed to confirm the delay and provide a more reliable and accurate measurement of its value. In the meanwhile, we sought to investigate coupled models of Rac1 and Cdc42 regulation that could quantitatively reproduce experimentally observed dynamics and proposed alternative signaling models that can explain the simultaneous (in the case of breast cancer cells) or delayed (in the case of MEFs) activation of both GTPases. In the first model, Rac1 and Cdc42 form a bidirectionally coupled system, where Cdc42 activity forms a polarized pattern that defines regions of high protrusive activity, while Rac1 activity drives the protrusion-retraction cycle with feedback to Cdc42. In the second model, both Rac1 and Cdc42 respond to the same upstream regulator. We hypothesize that such a regulator can work through the PI3K signaling pathway and activate Rac1 and Cdc42 activity through phosphoinositide regulation. In the third model, in the presence of a common upstream regulator, we introduced the crosstalk between two GTPases in the form of positive feedback from Cdc42 to Rac1. We showed that such crosstalk could synchronize Cdc42 and Rac1 dynamics and compensate for the delayed activation. Our computational framework can be used as a platform to study other types of cellular morphodynamics and their regulation by other signaling pathways of interest. The output of the model matches the discretization and resolution of experimental images, which makes it straightforward to compare simulation results with microscopy data using the same methods of analysis. We envision that by reaching a quantitative agreement between the data and our simulations of the complex interplay between GTPase activity and cell edge dynamics, this work will stimulate further development of integrative models to study the multiscale regulation of morphodynamics in different cell types and under different conditions.

## Materials and Methods

### Reaction-diffusion models of GTPase signaling

The general reaction-diffusion formulation of the spatial and temporal activity of Rho-GTPases in active and inactive forms can be presented as

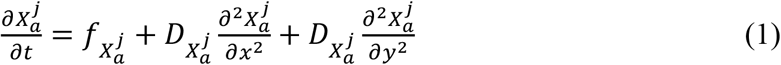

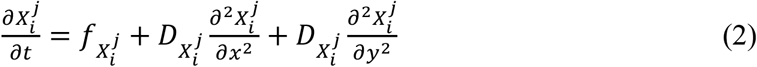

where *X*_*a*_ and *X*_*i*_ are two components representing concentration of protein *X* in its active and inactive forms, respectively, *j* is the numerical index, *D* is the diffusion coefficient, and *f* is the reaction term for each component.

For solving a 1D version of these equations, we use the built-in MATLAB finite element solver, *pdepe*, with the size of the simulation domain equal to 4 a.u. and the duration of the simulation domain equal to 10^3^ a.u. (for modeling of static GTPase activity patch in the two-component system) or 5 ⋅ 10^5^ a.u. (for modeling of dynamic GTPase activity in the four-component system). We use this solver for both homogeneous (spatially uniform GTPase activity) and heterogeneous (local spike of GTPase activity) initial conditions.

As a minimal 1D mass-conserved RD (MCRD) model of a single GTPase, we followed the previously implemented [39, 48, 49] form of equations:

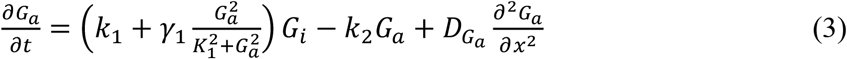

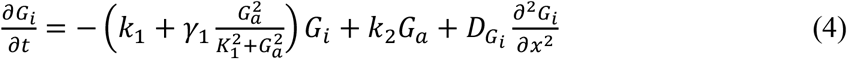

where *G*_*a*_ and *G*_*i*_ are the concentrations of active and inactive forms of the GTPase, respectively. The numerical values of the parameters (in arbitrary units) used in this work are *k*_1_ = 0.005, *γ*_1_ ∈ [0,6], *K*_1_ ∈ [0, 10], *k*_2_ = 0.1, *G*_*t*.*c*._ = 1 (the total GTPase concentration), 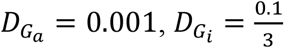. As the homogeneous initial conditions, we used the values 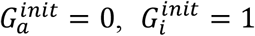. As the heterogeneous initial conditions, we used: 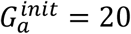 for *x* ∈ [0.0,1] 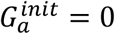 for *x* ∈ [0.1, 4], 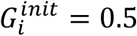 for *x* ∈ [0, 4]. We also added a small random noise to the initial conditions to create a perturbation to the unstable homogeneous state: 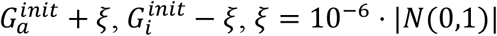, where *N*(0,1) is a normally distributed random variable with a mean equal to zero and a standard deviation equal to one.

As a 1D, four-component model for GTPase activity with negative feedback, we implemented the following equations:

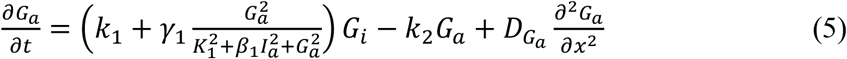

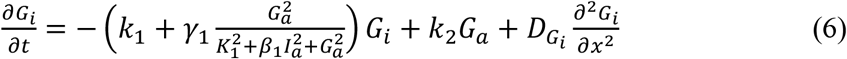

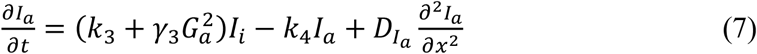

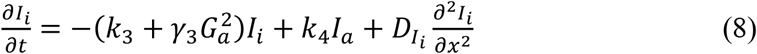

Here *I*_*a*_ and *I*_*i*_ are the concentrations of an inhibitor in its active and inactive forms. We assume that the inhibitor increases the deactivation rate of a GEF upregulating GTPase activity (see

### Supplemental Text

for details). The numerical values of the parameters (in arbitrary units) are *k*_1_ = 0.005, *γ*_1_ = 2, *K*_1_ = 1.4 (for oscillatory dynamics), *K*_1_ = 2.1 (for excitable dynamics), *β*_1_ = 0.5, *k*_2_ = 0.1, *k*_3_ = 10^−5^, *γ*_3_ = 0.2, *k*_4_ = 10^−2^, *G*_*t*.*c*._ = 1 (the total GTPase concentration), *I* = 3 (the total inhibitor concentration), 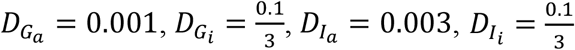. For the analysis of excitable dynamics, we used the following initial conditions: 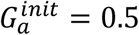 for *x* ∈ [1.4, 1.6], 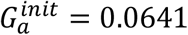 for *x* ∉ [1.4, 1.6] 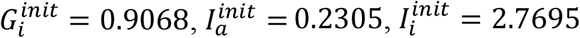. For the analysis of oscillatory dynamics, we used: 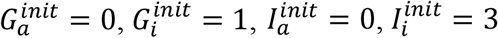. Similarly to the case of the one-component model, we added small perturbation to the initial conditions: 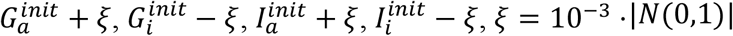.

To obtain the numerical solution for 2D models, we applied the forward Euler method with split reaction and diffusion operators with the time step Δ*t* = 0.001 and spatial grid step Δ*x* = 0.02. The 2D version of the RD equations also included a noise term to account for intrinsic cellular noise and stochastic switching of GTPases and their regulators between active and inactive forms:

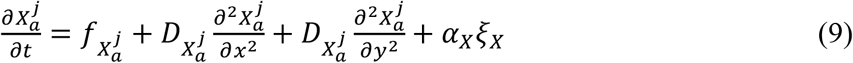

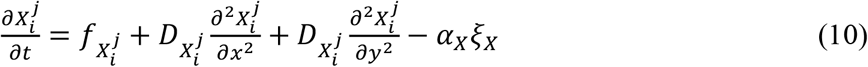

Here *α*_*X*_ is the amplitude of the noise and *ξ*_*X*_ is a Gaussian random variable with a mean of zero and a standard deviation of one. The noise terms in the first and second equations have the opposite signs and represent stochastic switching between the active and inactive forms of component *X*. To ensure mass conservation and positivity of the solution, we applied additional checks at each iteration, as described in [44]. We solved the equations in an arbitrarily shaped simulation domain representing cell shape with no-flux boundary conditions. To this end, we used the discrete Laplace operator (based on a five-point stencil) with components representing fluxes across the edge of the simulation domain set to zero. For the details of this implementation, see [44]. As a static simulation domain (i.e., in the case of a non-moving cell), we used a square domain of 200×200 grid points (4×4 a.u). For simulations with a dynamic cell, we used either a circular initial simulation domain of a radius equal to 80 pixels or a cell mask extracted from the experimental data.

The 2D model for GTPase activity during cell ruffling was implemented in the following form:

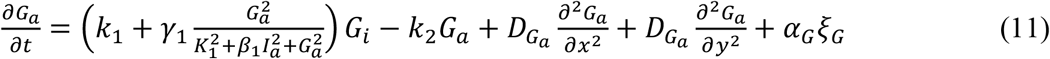

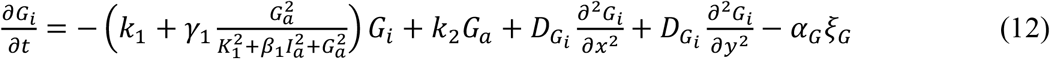

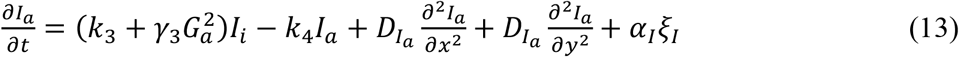

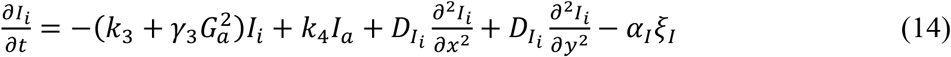

The values of kinetic parameters were the same as in the 1D model except for the *K*_1_ parameter, which was set differently at the border of the simulation domain and elsewhere. Based on experimental evidence [12, 50-52], we assumed an increased GTPase activity at the narrow (1-pixel) band on the very edge of the cell. Thus, at the boundary nodes of the simulation domain, we set 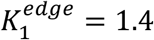 and for the rest of the domain, we set 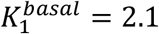. In all figures of the Results section, the values of noise were set to *α*_*G*_ = *α*_*I*_ = 5. The initial conditions were set to 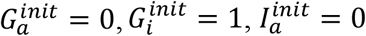, and 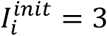.

To investigate the response of the 2D model of GTPase activity during cell ruffling to different values of noise, we performed a series of simulations with different values of noise amplitude (**Supplemental Figure 1**). We also applied textural analysis for the quantification of patterns (see [44] for this method of analysis). With small values of noise, patterns were not formed because diffusion fluxes had sufficiently high values to prevent pattern formation at the edge of the domain. For an increased noise (*α* = 5), patterns started to form. However, for even higher values of noise, patterns become suppressed by the overly strong perturbation. Based on these results, we have chosen the value of *α* = 5 for our simulations.

As a next step in model development, we implemented a 2D model of coupled Cdc42 and Rac1, where Rac1 activity is induced by Cdc42:

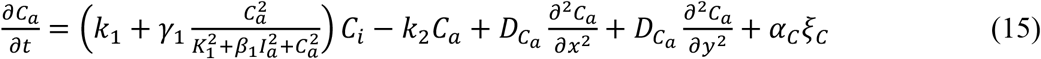

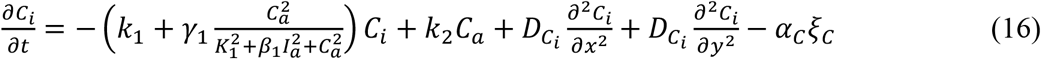

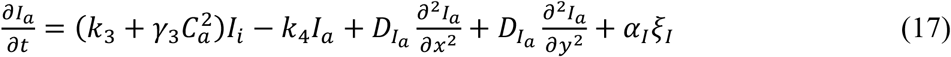

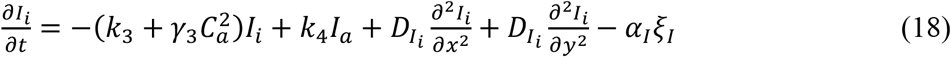

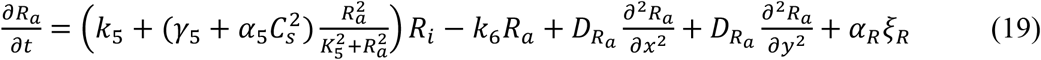

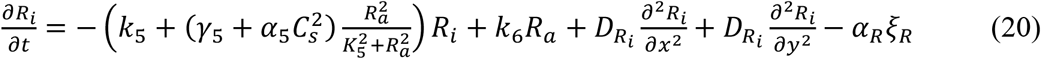

Here, the numerical values of the parameters (in arbitrary units) are 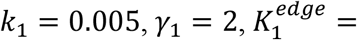 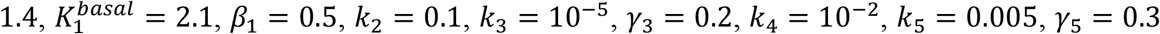 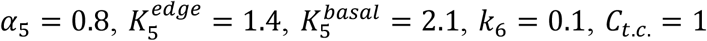 (total Cdc42 concentration), *I*_*t*.*c*._ = 3 (total inhibitor concentration), *R*_*t*.*c*_ = 1 (total Rac1 concentration), 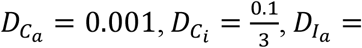 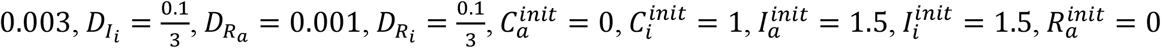, 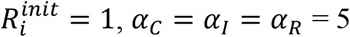.

Next, we explored a 2D model of Cdc42 and Rac1 with a bidirectional coupling. In this model, Cdc42 controls cell polarization, and Rac1 drives protrusion-retraction dynamics:

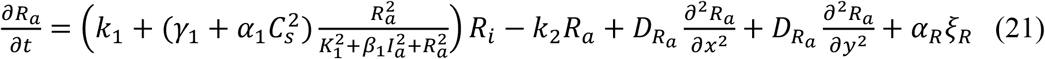

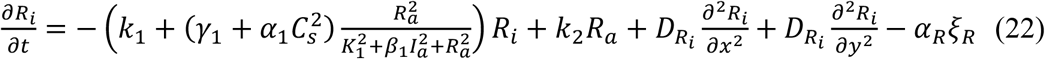

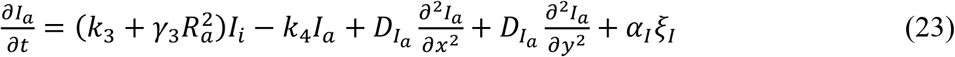

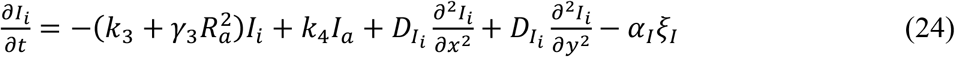

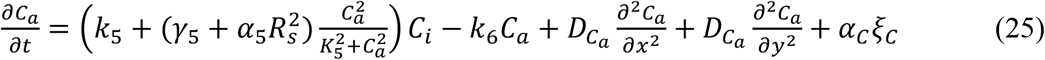

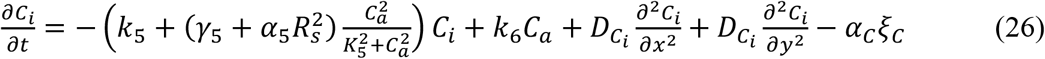

Here the numerical values of the parameters (in arbitrary units) are: 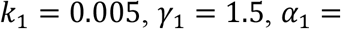 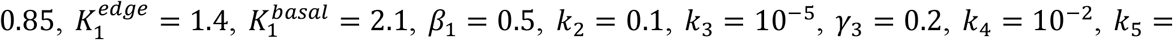 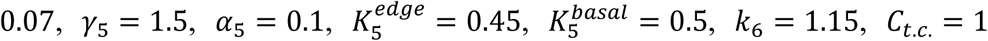 (total Cdc42 concentration), *I*_*t*.*c*._ = 3 (total inhibitor concentration), *R*_*t*.*c*._ = 1 (total Rac1 concentration), 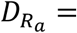 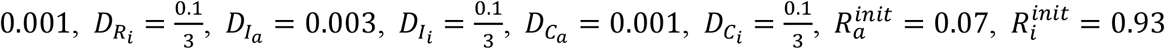, 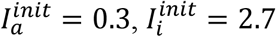. For Cdc42 concentration, we used heterogeneous initial conditions with a local activation stimulus, 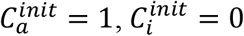, and the otherwise inactive state 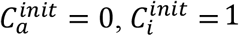. The values of noise amplitude were: *α*_*R*_ = *α*_*I*_ = *α*_*C*_ = 5.

A 2D model with both Cdc42 and Rac1 regulated by an upstream signaling motif was implemented in the following form:

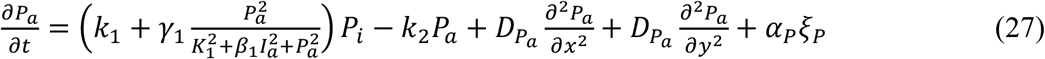

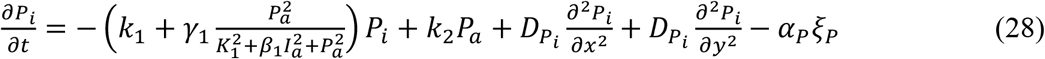

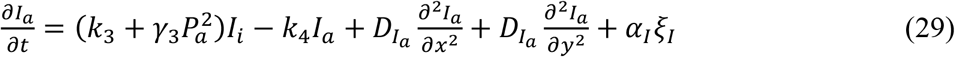

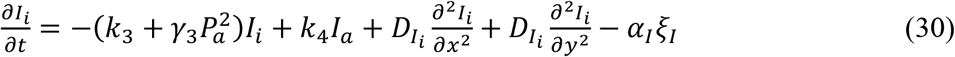

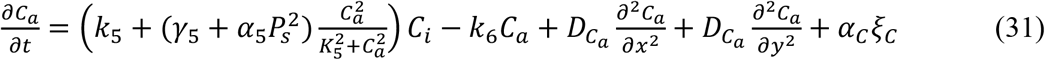

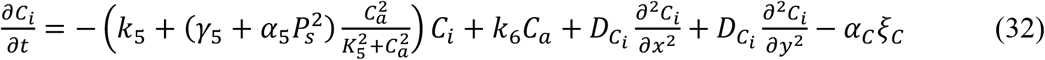

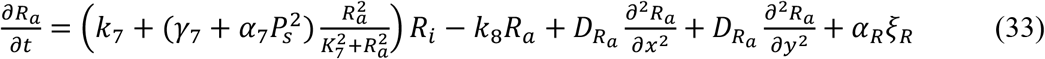

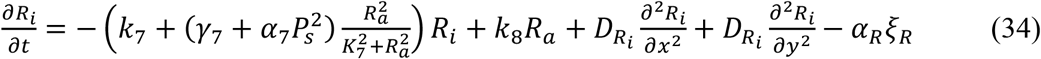

The numerical values of the parameters (in arbitrary units) are 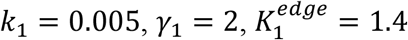, 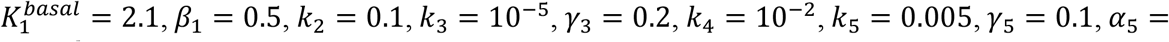 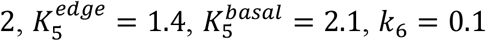. For the case of simultaneous Rac1 and Cdc42 activity, we used the following parameters in the Rac1 motif: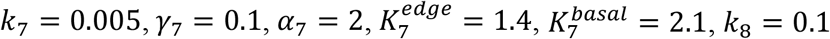. For the case of a delayed Rac1 activation, we used: 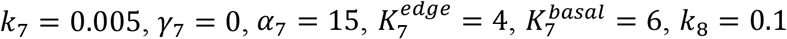. In all simulations of this model, we used: *P*_*t*.*c*._ = 1 (total concentration of the upstream component), *I*_*t*.*c*._ = 3 (total inhibitor concentration), *C*_*t*.*c*._ = 1 (total Cdc42 concentration), *R*_*t*.*c*_ = 1 (total Rac1 concentration), 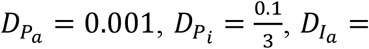 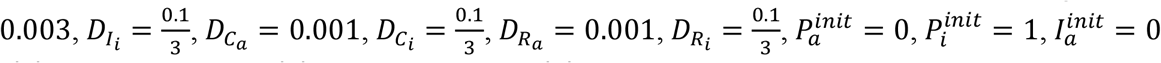, 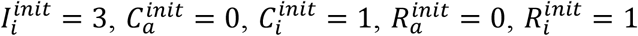. The values of noise amplitude were *α* = *α*_*I*_ = *α*_*C*_ = *α*_*R*_ = 5.

Finally, a unified 2D model with Cdc42 and Rac1 regulated by both feedback from Cdc42 to Rac1 and an upstream signaling motif was implemented in the following form:

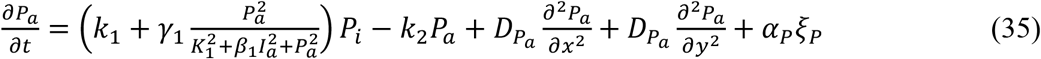

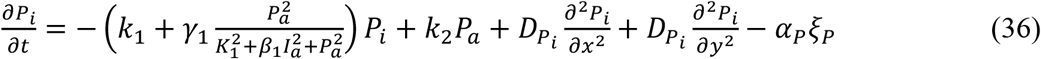

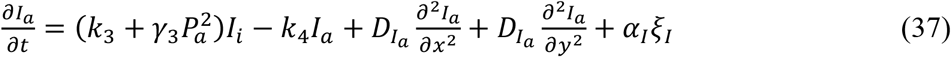

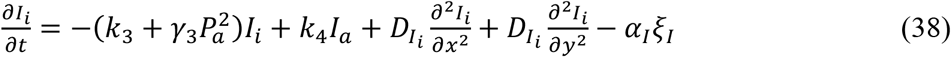

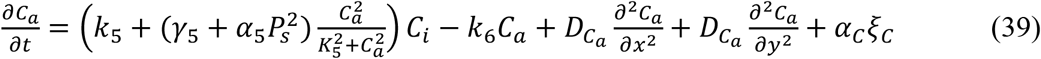

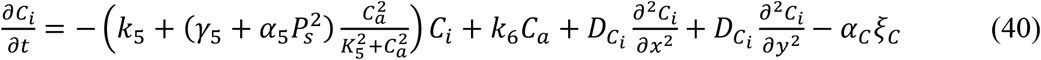

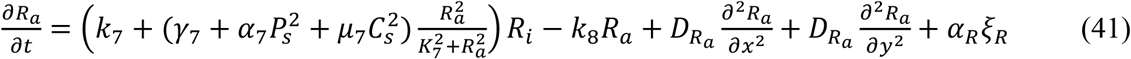

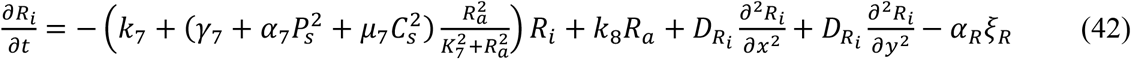

The numerical values of the parameters (in arbitrary units) are 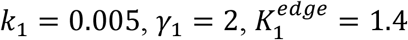, 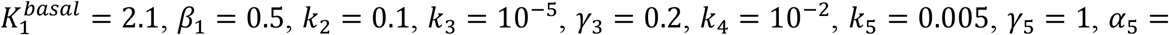 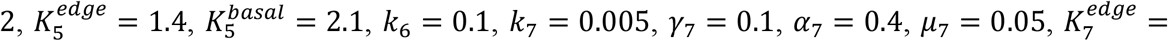 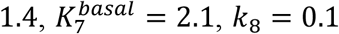. For the case of a delayed Rac1 activation, we used: *μ*_7_ = 1. In all simulations of this model, we used: *P*_*t*.*c*._ = 1 (total concentration of the upstream component), *I*_*t*.*c*._ = 3 (total inhibitor concentration), *C*_*t*.*c*._ = 1 (total Cdc42 concentration), *R*_*t*.*c*._ = 1 (total Rac1 concentration), 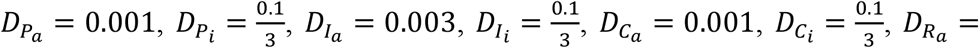 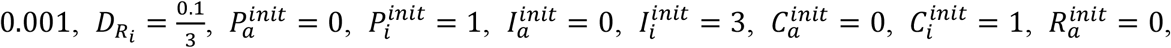 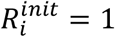. The values of noise amplitude were *α*_*P*_ = *α*_*I*_ = *α*_*C*_ = *α*_*R*_ = 5.

### A computational method for coupling reaction-diffusion system of equations with cellular morphodynamics

The rationale and general outline for our modeling approach are provided in the Results section. Here, we describe the technical details of its implementation.

The probabilities of local protrusion and retraction were defined as the multiplication of several contributing factors. The ‘geometry factor’ accounts for the local influence of membrane curvature. It is calculated for each pixel depending on the values of the cell mask in the 8-connected neighborhood of that pixel. The formulas below ensure that the regions with large positive curvature have a lower probability of further protruding and a higher probability of retracting. In contrast, the regions with large negative curvature have a higher probability of protruding and a lower probability of retracting. The geometry factors for protrusion and retraction probabilities are, respectively:

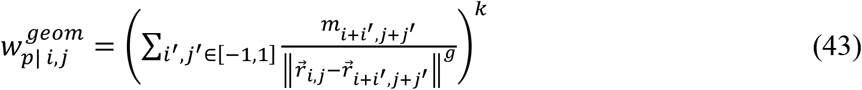

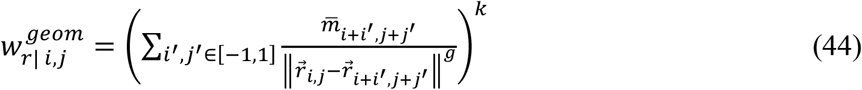

where *m*_*i,j*_ represents the value of a binary cell mask in position 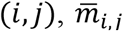 is the value of the inversed mask 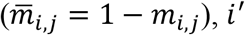 and *j*^′^ are pixel locations in the 8-connected neighborhood of pixel (*i, j*), 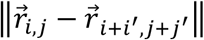 is the Euclidian distance between positions (*i, j*) and (*i* + *i*^′^, *j* + *j*^′^), *g* defines the sensitivity of the geometry factor to the curvature, and *k* defines the relative contribution of this factor to the overall probability (i.e., higher values of *k* lead to a smoother appearance of the cell outline). Hereon, the indexes ‘*p*’ and ‘*r*’ stand for protrusion and retraction, respectively.

The ‘volume factor’ controls the conservancy of cell size with a step-like dependence, ensuring that the increase of cell volume decreases the probability of further protrusion and increases the probability of retraction. Correspondingly, the decrease in cell volume has the opposite effect.

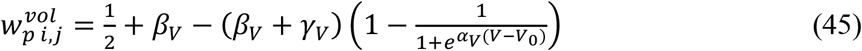

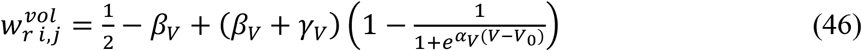

Here *V* and *V*_0_ represent the current and initial volumes of the cell, and *α*_*V*_ regulates the sharpness of the step function. When the volume increases significantly compared to 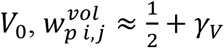 and 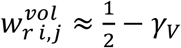, so that *γ*_*V*_ is the deviation of protrusion and retraction probabilities from the equal 0.5 value. In contrast, when the volume decreases considerably with respect to 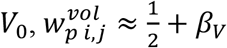, and 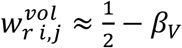, so that the deviation of protrusion and retraction probabilities from 0.5 is defined by the value of *β*_*V*_. Thus, parameters *β*_*V*_ and *γ*_*V*_ together define the sensitivity of cell dynamics to the deviation of its volume from a constant value.

The ‘actin factor’ is introduced similarly to the volume factor but with opposite signs in the step-like function to ensure that higher rates of actin polymerization lead to protrusion and lower rates lead to retraction:

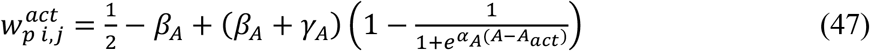

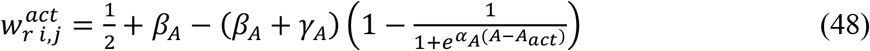

Here *A* represents the effects of actin polymerization on the protrusion and retraction probabilities. Such effects can be influenced by the concentration of active GTPase molecules or by the kinetic rate of GTPase activation (the need for this distinction is described in the Results section). *A*_*act*_ defines the value of *A* at the inflection point of the step-like function. In all reported results, we assumed *β*_*A*_ = 0, *γ*_*A*_ > 0, which implies that weak actin regulation does not create a relative shift in the protrusion and retraction probabilities, while upregulation (*A* > *A*_*act*_) leads to an increased protrusion rate and a decreased retraction rate.

In our models, changes in cell shape result from the addition (protrusion event) or removal (retraction event) of a pixel at the edge of the cell mask. Such events are determined by calculating the probabilities of protrusion and retraction for each foreground and background pixel along the outline of the cell mask. To calculate the actin factor for a background pixel, we extrapolate the values of concentrations (or kinetic rates) from the RD model by averaging the values in the pixels of the cell mask within the 8-connected neighborhood of this background pixel:

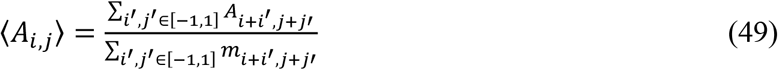

Based on these probabilities, a random number generator selects a set of added and removed pixels. During the update of the cell mask (both for protrusion and retraction steps), we apply an additional automated filtering step that guarantees the unity of the simulation domain: i.e., the algorithm prohibits events that lead to spur (diagonal 4-disconnected) pixels or the formation of hollow or 4-disconnected parts of cell mask. To guarantee mass-conservation of the signaling components, we subtract the total molecular mass in the added pixels evenly from all pixels in the new mask. Because the number of added pixels is always a small fraction of all pixels in the simulation domain, the subtracted value is relatively small. To ensure the positivity of concentrations, we also check that the subtraction is not applied to pixels with concentrations less than the subtracted value. With this approach, the increase in the area of the simulation domain leads to a decrease in the total concentration but not the total mass of the RD components, as it was also implemented in other studies [53]. Similarly, for retraction events, the total molecular mass of the components in the RD model from the removed pixels is evenly distributed across the whole new mask. With this approach, a decrease in the size of the simulation domain leads to an increase in concentration values while the mass is still conserved. After one protrusion and one retraction event, we run a sequence of 50 iterations of the RD model with the forward Euler method in the updated cell mask (as described above). The state of the system was saved after every 1000 iterations of the RD model, which is the 1 a.u. of the simulation time.

The values of parameters associated with geometry and volume factors were the same in all models that we used in this work: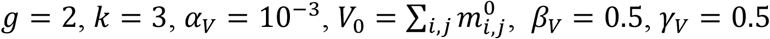. For the actin factor, we adjusted parameters separately for each model because of the different kinetic parameters in RD equations. The parameters were adjusted to (1) select the threshold for activation of the actin factor (*A*_*act*_), (2) define the sharpness of the sigmoid function (*α*_*A*_) that matches the timescale of changes in the concentration of GTPase or the kinetic rate of its activation, and (3) match the protrusion rate by adjusting *γ*_*A*_ parameter.

In all simulations presented in this work, we used the value *β*_*A*_ = 0. The values of other parameters related to the actin factor are shown in the following **Table 1**:

**Table 1.**
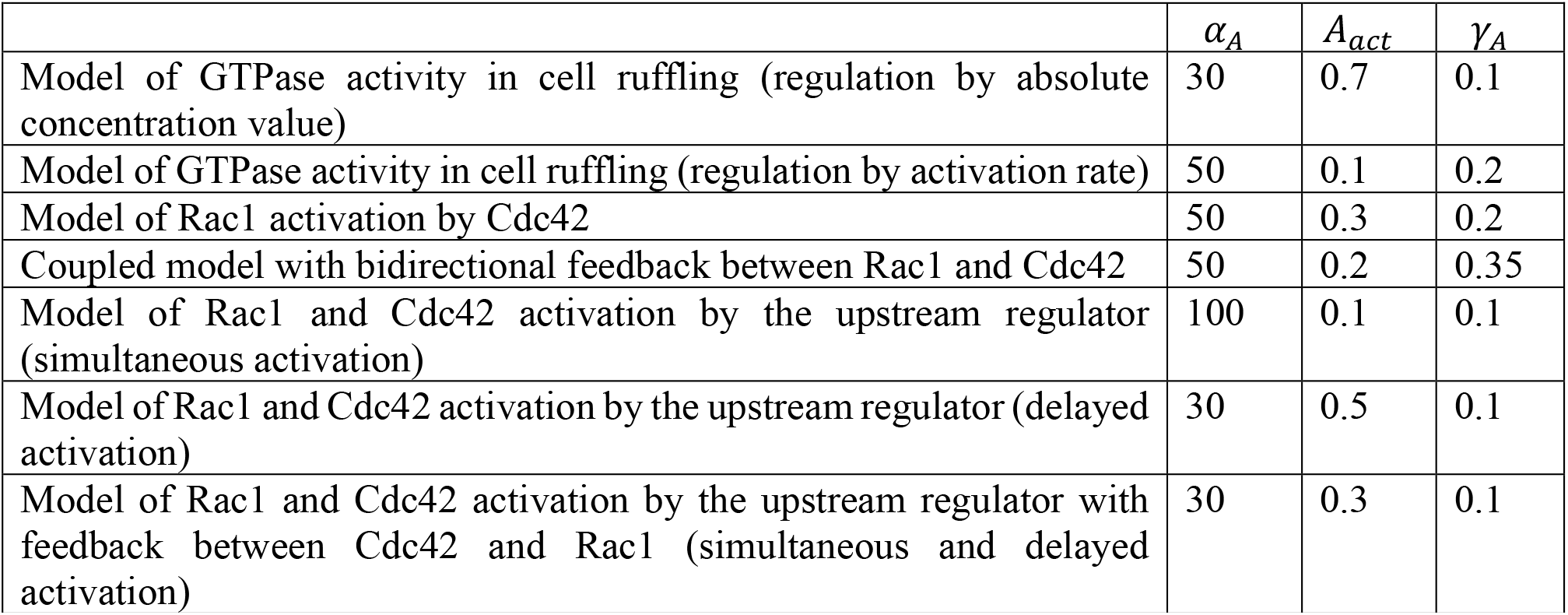
Parameter choices for cell morphodynamic models. The other parameters were the same for all models and provided in the text of the Method section.

| | $\alpha_A$ | $A_{act}$ | $\gamma_A$ |
| --- | --- | --- | --- |
| Model of GTPase activity in cell ruffling (regulation by absolute concentration value) | 30 | 0.7 | 0.1 |
| Model of GTPase activity in cell ruffling (regulation by activation rate) | 50 | 0.1 | 0.2 |
| Model of Rac1 activation by Cdc42 | 50 | 0.3 | 0.2 |
| Coupled model with bidirectional feedback between Rac1 and Cdc42 | 50 | 0.2 | 0.35 |
| Model of Rac1 and Cdc42 activation by the upstream regulator (simultaneous activation) | 100 | 0.1 | 0.1 |
| Model of Rac1 and Cdc42 activation by the upstream regulator (delayed activation) | 30 | 0.5 | 0.1 |
| Model of Rac1 and Cdc42 activation by the upstream regulator with feedback between Cdc42 and Rac1 (simultaneous and delayed activation) | 30 | 0.3 | 0.1 |

### Image analysis pipeline for coupled analysis of cell edge velocity and biosensor signal

A conceptual overview of our image analysis pipeline is provided in the Results section. Here, we describe the technical details of its implementation.

As input for our analysis, we used experimental FRET biosensor data with the ratio signal [17, 47]. For plotting, we adjusted the gray-scale intensity limits to 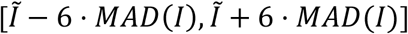, where 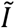 is the median value of the biosensor signal across the whole time series data and *MAD*(*I*) is a median absolute deviation. We also disregarded outliers in the histogram of the biosensor signal if the number of pixels outside the interval [*I* − 5, *I* + 5] was less than 20. After that, the signal was scaled as (*I* − min(*I*))/(max(*I*) − min(*I*)).

Using our mid-contour approach described in the main text, we build a large set of trajectories along which each individual point of the cell outline moves over the course of the whole time-lapse recording. Such trajectories represent local directions of fast protrusion/retraction cycles, while the curving of these trajectories reflects the slow change of the overall cell shape over multiple protrusion/retraction events.

To analyze the biosensor signal in the neighborhood of local edge velocity maxima, we first filtered out trajectories with absolute values of velocity larger than 20 standard deviations (which usually represent cell segmentation artifacts). We identified regions of kymographs with high values of velocity using the criteria: 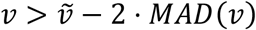 and filtered out regions of a size smaller than 10 pixels. The points of local velocity maxima were found in the identified regions. The values of cell edge velocity and the biosensor signal were analyzed in the time interval ±20 frames with respect to the time of the velocity peaks. We then computed the mean values at each time point (along all trajectories) and computed 97.5% confidence intervals. The temporal derivative of the biosensor signal was computed based on the difference from the previous time frame. The coordinate of the membrane was computed as an integral of velocity starting from the first frame in the time interval.

For the analysis of multiplexed Rac1 and Cdc42 data, we processed each GTPase channel separately and combined the results in the time intervals around the time of velocity peaks. Data were averaged over all 6 cells in each dataset.

The same pipeline for the analysis of GTPase concentration was applied to quantify our simulation results. The simulated time series was recorded with a 10 a.u. step.

To match the time and space units between simulations and experimental data, we used temporal and spatial autocorrelation plots computed based on velocity kymographs (**Supplemental Figure 2**). Based on this analysis, 1a.u. of simulation time is equal to 0.65s and 1a.u. of spatial dimension is equal to 10.127 microns.

## Results

### Coupling reaction-diffusion models of GTPase regulation and cell morphodynamics

#### 1. Modeling cell morphodynamics

We build a model of a moving cell as a sequence of stochastic protrusion-retraction events. The probability of each protrusion and retraction event is modulated by GTPase Rac1 activity through actin polymerization. For direct comparison of the model output with experimental microscopy data, we performed the simulation on a square grid with the resolution matching the imaging data. We represent a cell of an arbitrary shape as a binary 4-connected object on a square grid (also referred to as the cell mask *M*). The cell shape is updated stochastically by adding pixels (protrusion) and removing pixels (retraction) at the cell edge (**Fig. 1A**). Conceptually, such an approach is similar to the Cellular Potts Model (CPM) [54]. However, our implementation has some major differences. Specifically, in our model, we separate protrusion and retraction events, which allows us to distinguish between the regulations of these two processes and model them differently if needed. We also do not introduce a global Hamiltonian to define the probabilities of cell shape changes because such formulation is somewhat abstract and cannot be easily related to cell morphodynamics. Instead, we aim to build a model that can be related to biochemical regulation and be interpreted from the biomechanical perspective more directly. Thus, we defined the probabilities of protrusion and retraction with three contributing factors: local geometry (local curvature of the cell membrane), overall cell volume (change in the size of the simulation domain), and local actin polymerization (controlled by the activity of GTPases in the RD model) (**Fig. 1B**).

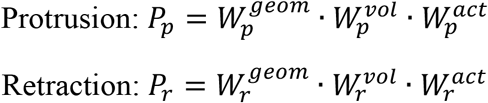

**Figure 1.**
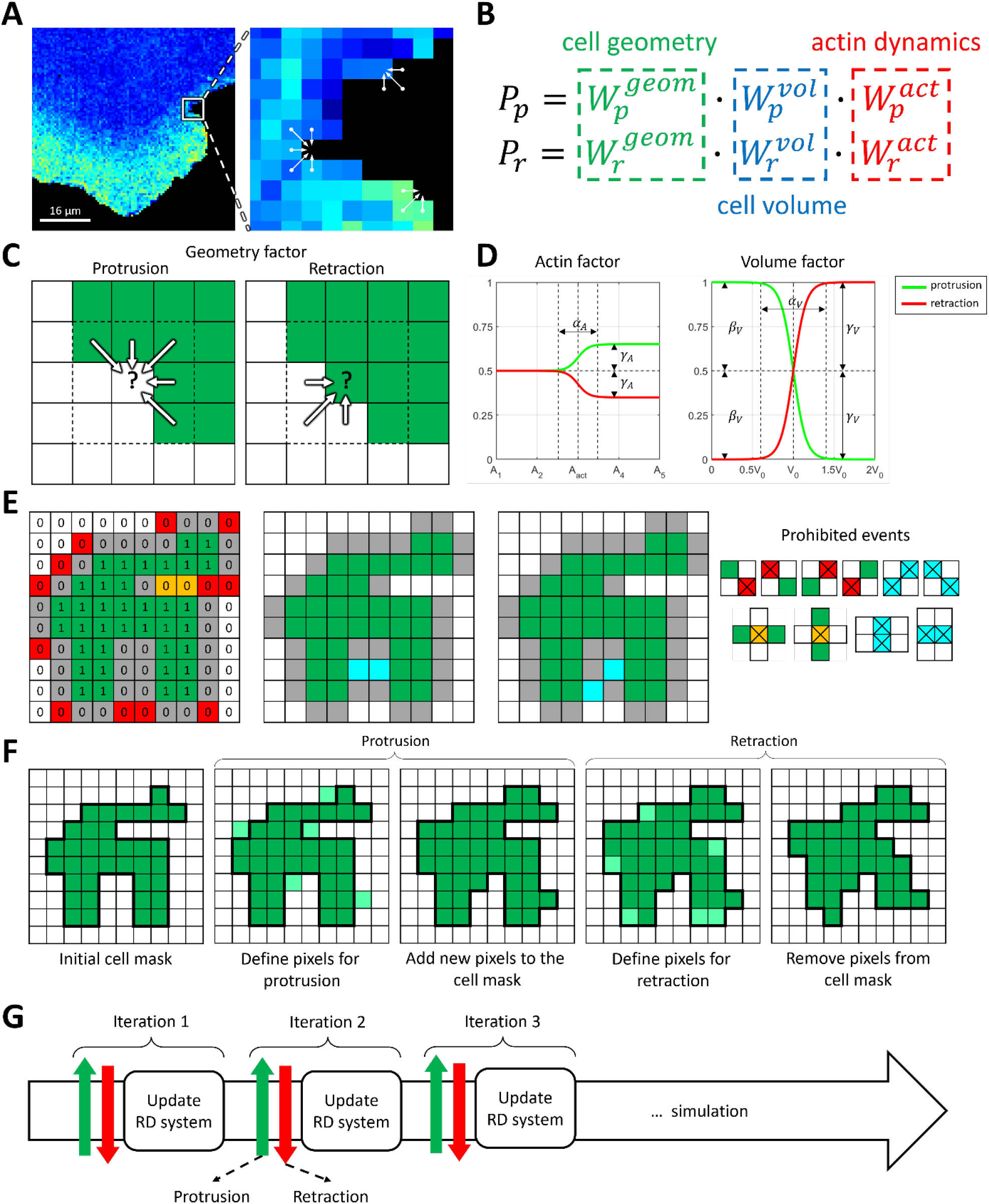
Integrative model of cell morphodynamics. **A**. Cell is represented as a binary 4-connected object on a square grid. The change of cell shape is represented as a series of protrusion/retraction events. **B**. Probabilities of protrusion and retraction depends on three factors: geometry factor (green), volume factor (blue) and actin factor (red). **C**. Geometry factor is defined for each pixel at the outline of the cell and depends on the values of cell mask (green) in the neighborhood. Convex regions of cell mask with high curvature are less likely to protrude and more likely to retract. Concave regions of the cell mask with negative curvature are more likely to protrude and less likely to retract. **D**. Actin and volume factors have a form of sigmoid functions and represent deviation from 0.5 value depending on the regulator of actin or cell volume. Parameters include the threshold of activation (*A*_*act*_, *V*_0_), sharpness of the response (*α*_*A*_, *α*_*V*_), deviation from 0.5 for increased regulation (*γ*_*A*_, *γ*_*V*_), and decreased regulation (*β*_*V*_). **E**. Protrusion events take place at the outline (gray 0s) of the cell mask (green 1s). To maintain the unity of simulation domain we prohibit diagonal events (red 0s). Events that lead to the formation of bubbles are also prohibited (orange 0s). Simultaneous updates of the pixels at cell outline can lead to the formation of bubbles or diagonal events (blue). Such events are processed separately so that only one of them takes place. During retraction the logic is reversed: 0s represent cell mask and 1s represent empty space. **F**. During cell shape update protrusion event is followed by retraction event. **G**. Cell dynamics is represented as an iterative process where protrusion-retraction cycle is followed by the update of intracellular signaling (reaction-diffusion, RD model).

For each of these factors, we parametrize the probability of protrusion and retraction so that they contribute to the membrane velocity depending on biological and physical rationale. Specifically, the geometry factor describes the response to local membrane curvature (**Fig. 1C**) so that convex regions of the cell mask with high positive curvature are less likely to protrude and more likely to retract. In contrast, concave regions of the cell mask are more likely to protrude and less likely to retract. This setup is well suited for modeling broad lamellipodia-like protrusions, such as during cell ruffling. For modeling filopodia-like protrusion or other high-curvature cell shapes, the setup could be modified accordingly. The volume factor accounts for the effects of cell size changes, which may happen on a larger time scale than a single protrusion/retraction event. In our implementation, the increase of the cell size with respect to its initial size leads to decreased protrusion and increased retraction probabilities and vice versa (**Fig. 1D**). As a result, cell size remains close to the initial value with some stochastic deviations. The extent of such deviations can be regulated by the parameters of the model (see Methods). Finally, the actin factor describes the regulation of cell morphodynamics by GTPase signaling pathways that induce actin polymerization driving cell edge movement. We assume that in the absence of GTPase activity, the regulation is neutral, i.e., this factor does not contribute to the relative change in the probabilities of protrusion and retraction (**Fig. 1D**). In other words, when the concentration of the active form of GTPase is below the specified threshold, the actin factor is equal to 0.5 for both the protrusion and retraction probabilities. The upregulation of GTPase activity leads to an increased protrusion rate and decreased retraction rate. In such a setup, the cell protrusion (when GTPase is locally activated) is followed by a relaxation phase (when GTPase is deactivated) when the cell membrane retracts based on the other probability factors. The functional forms of each factor are described in the Methods section.

Based on the described probabilities, the overall dynamics of cell shape is calculated as a sequence of protrusion/retraction events using the Monte-Carlo algorithm (**Fig. 1E-G**). During the protrusion step, new pixels are added to the cell edge (i.e., some of the background pixels at the very edge of the cell mask become foreground pixels). During the retraction step, pixels are removed from the cell edge (i.e., some of the pixels forming the cell outline become background pixels). To maintain the 4-connectedness, unity of the cell mask, and the absence of holes, our algorithm automatically prohibits pixel addition/removal that would lead to such distortions.

#### 2. Spatiotemporal model of GTPase activity during cell ruffling

FRET biosensor data [17, 18] shows that during cell membrane ruffling, Rac1 and Cdc42 activities correlate with membrane velocity and form transient localized patches of activity close to the cell edge. To represent such dynamics in simulations, we used a MCRD model with autocatalytic feedback as a core signaling motif. Such a model can be interpreted as a coarse-grained approximation of a more complex signaling motif with positive feedback through the activation of GTPase effectors (e.g., GEFs), which in turn increases the activation of GTPase (**Fig. 2A**). Mathematically, converting the extended model into the coarse-grained one (see **Supplemental Text**) leads to the dependence of parameter *γ*_1_ (the maximum of the resulting Hill function) on the total concentration of GTPase effectors, such as GEF, and parameter *K*_1_ (the threshold of activation) on the rate of effector deactivation. Depending on its parameters, the MCRD model can operate in the excitable regime or in the Turing-unstable regime. In the excitable regime, a finite stimulus is needed to induce the formation of an activation patch, and in the Turing-unstable regime, the homogeneous state is unstable, and no stimulus is needed to induce GTPase activation (**Fig. 2B,C**). Outside of these regimes, the system is in an inactive state when no patterns are formed with or without a stimulus. When an activity patch is formed in either the excitable or Turing-unstable regime, it can be deactivated by a regulator that moves the system to the inactive regime. We visualized this behavior with the phase space portrait for parameters *γ*_1_ and *K*_1_ (**Fig. 2B,C**). Thus, in order to model a system with the transient formation of an activity patch, we coupled the core signaling motif and a regulator that modulates the positive feedback in the system (e.g., negatively regulates the activation of GEFs). This way, active GTPase increases the activation of a regulator that deactivates the GTPase by increasing the threshold of the autocatalytic activation (*K*_1_ parameter, **Fig. 2D,E**). This coupled system can be excitable or oscillatory. In the excitable state, the activity is induced by a stimulus and decays due to the negative feedback from the regulator, and in the oscillatory state, the core motif switches between Turing-unstable and inactive states due to the negative feedback from the regulator (**Fig. 2F-H**). Previously, it was experimentally shown that in a narrow band right at the cell edge, GTPase GEF Asef remains active as the cell ruffles, while in the other areas, GEF activity varies during the protrusion/retraction cycle [50]. Such persistent GEF activation at the very edge of the cell may reflect the fact that many GTPase effectors contain the curvature-sensing BAR domain [51, 52, 55]. Thus, for the 2D implementation of our model, we assumed that the activity of GTPases is slightly higher at the boundary of the simulation domain than everywhere else. Furthermore, to achieve the spontaneous formation of activity patches as observed in cell ruffling, we chose parameter *K*_1_ so that at the boundary of the simulation domain, the system operates in the oscillatory regime, while inside the simulation domain, it is in the excitable state. Such a setup reproduces localized and transient activity of GTPases at the edge of the simulation domain, matching the experimentally observed dynamics during cell ruffling (**Fig. 2I**).

**Figure 2.**
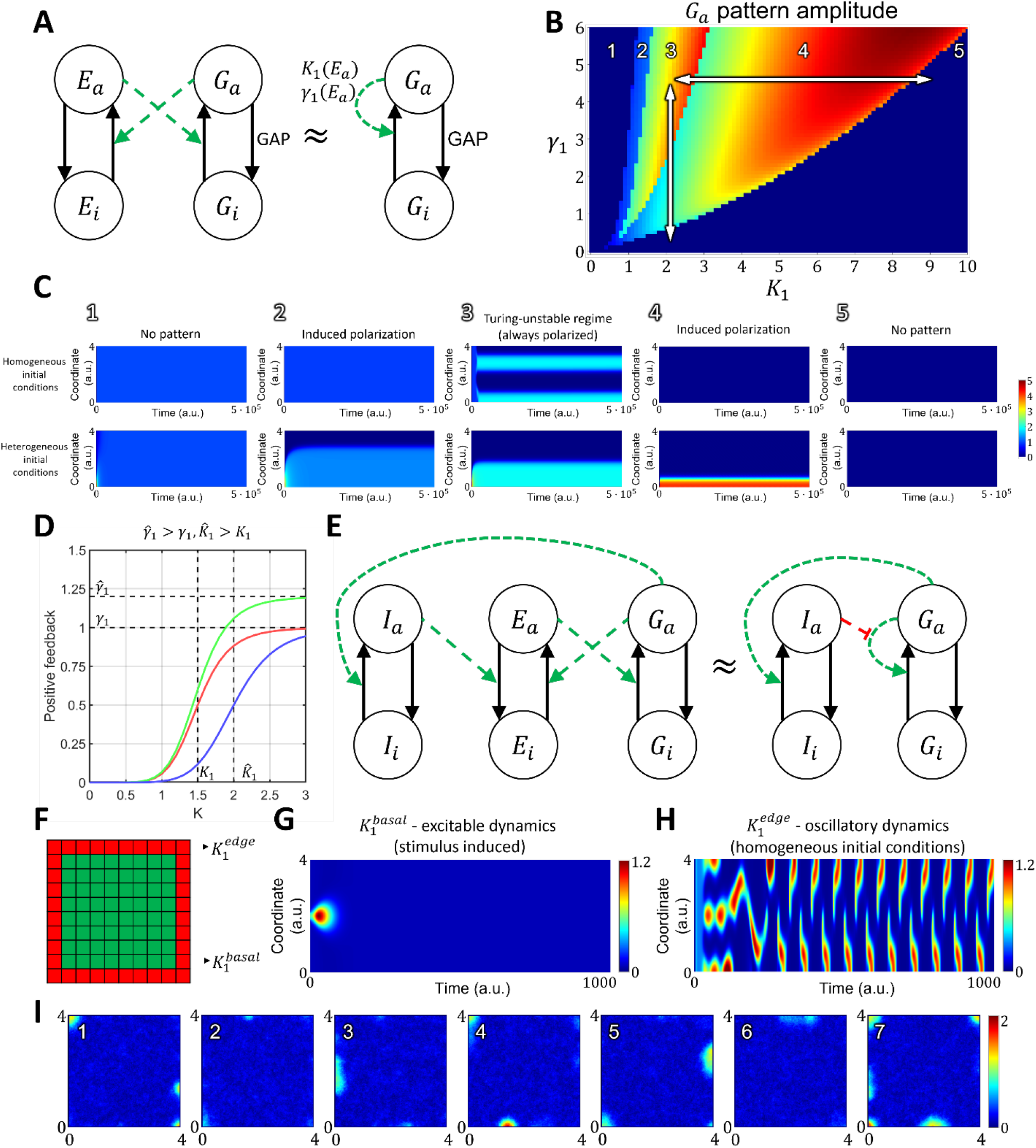
Reaction-diffusion model of GTPase activity in cell membrane ruffling. **A**. The two-component MCRD model for GTPase activity (*G*_*a*_ - active GTPase, *G*_*i*_ - inactive GTPase) can be interpreted as an approximation of an extended four-component, where active GTPase increases the activation rate of GEF (*E*_*a*_ - active GEF, *G*_*i*_ – inactive GEF) and active GEF, in turn, increases the activation rate of GTPase. **B, C**. In the phase space of *K*_1_ and *γ*_1_ parameters the MCRD model operates in the following regimes: 1. No pattern (stable, homogeneous, high-activity state, stimulus can’t induce polarization), 2. Stimulus-induced deactivation (stable, homogeneous, high-activity state, stimulus can induce polarization), 3. Turing-unstable regime (unstable homogeneous state, stimulus is not required to induce polarization, as it can be initiated by any small perturbation), 4. Stimulus-induced activation (stable, homogeneous, low-activity state, stimulus can induce polarization), 5. No pattern (stable, homogeneous, low-activity state, stimulus can’t induce polarization). **D**. The regulation of positive feedback in the MCRD changes the shape of the Hill function. An inhibitor can decrease *γ*_1_ parameter (magnitude of positive feedback) or increase *K*_1_ parameter (threshold of positive feedback activation). **E**. Signaling motif where active GTPase *G* activates inhibitor *I*, which in turn increases the rate of GEF deactivation. In the simplified model containing the MCRD motif, it is equivalent to the increase of *K*_1_ parameter. **F-H**. In 2D model we assumed an increased GTPase activity at the boundary of simulation domain (decreased 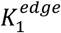 in comparison to 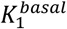), which corresponds to increased density of GEF proteins with curvature sensitive domain. 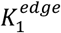 corresponds to the oscillatory dynamics, while 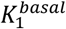 corresponds to the excitable dynamics. **I**. The GTPase dynamics in 2D model shows formation of transient activity patches at the boundary of the simulation domain which corresponds to the experimental observation, when GTPase (Rac1 and Cdc42) are activated at cell edge.

#### 3. A pipeline for the analysis of cell edge velocity and GTPase activity in experiments and simulations

For this work, we developed an image analysis pipeline allowing simultaneous analysis of cell edge velocity and biosensor signal during cell ruffling. This pipeline was motivated by the need to account for two timescales of cell movement: fast cell edge fluctuations (ruffling) and a slow change of the overall cell shape. To this end, we first processed the time-series data (**Fig. 3A**) by extracting cell masks (see **Methods**) and splitting the whole time series into 20-frame intervals. For each time interval, we found the outer contour, which represents the smallest region that encloses all cell masks in the time interval (i.e., the union of cell masks), and the inner contour, which represents the overlap of all cell masks in the time interval (i.e., the intersection of cell masks). The discretized representation of the contours was resampled with the 1-pixel distance between the consecutive contour points. Based on outer and inner contours, we use an iterative algorithm that converges to a mid-contour between the outer and inner ones. First, the algorithm computes normal lines for each point of the outer contour, finds their intersections with the inner contour, and sets the midpoints of these lines between the outer and inner contours as Contour 1. Next, the algorithm computes normal lines at each point of the inner contour, finds their intersections with the outer contour, and sets the midpoints as Contour 2. These two steps are repeated at the next iterations but with Contour 1 and Contour 2 instead of the outer and inner contours. The iterative process is interrupted when the average distance between the two new Contours 1 and 2 becomes smaller than a pre-defined tolerance (0.1 pixels, see **Methods**). Once the mid-contours are computed for each 20-frame time interval (**Fig. 3B**), we record the normal lines to these mid-contours and the intersection of each cell outline in this interval with the corresponding lines (**Fig. 3C**). Finally, we match the normal lines from consecutive time intervals based on the proximity of the last set of intersections in one interval and the first set of intersections in the next one. This way, we build piecewise linear trajectories, along which the points of the cell outline move over time (**Fig. 3D**).

**Figure 3.**
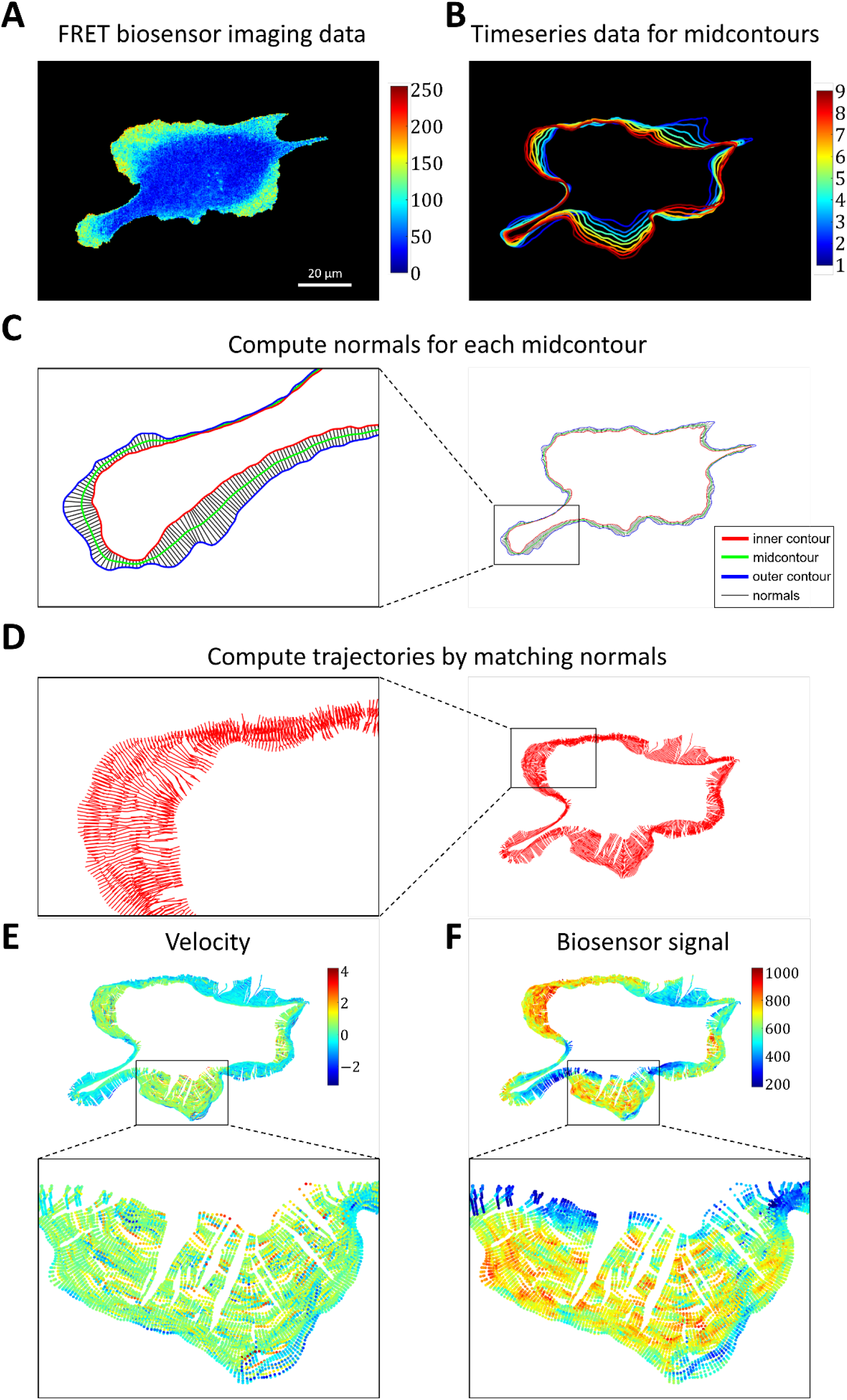
Image analysis pipeline for simultaneous analysis of cell edge velocity and biosensor signal. **A**. The input of the pipeline is time-series data of FRET biosensor data in cells. **B**. The time-series data was split into time intervals of 20 frames and for each time interval a midcontour was computed. **C**. For each computed midcontour the normals were computed and intersections with the inner and outer contours were found. **D**. Cell membrane trajectories were computed by matching the closest normals. **E, F**. Cell edge velocity and biosensor signal were tracked along the computed trajectories at the points of intersection with cell edge.

For each point of each trajectory, we compute the velocity (based on position change along the trajectory) and biosensor signal (by averaging the values in the neighborhoods of the points) (**Fig. 3E, F**). Using this method, we represent cell velocity and biosensor signal as kymographs, with the x-axis showing the numeric index of the trajectories and the y-axis showing time.

Finally, we use these kymographs to analyze the relationship between GTPase activity and dynamics of cell edge motion near the time and location of the velocity peaks (see **Methods, Fig. 4A-C**).

**Figure 4.**
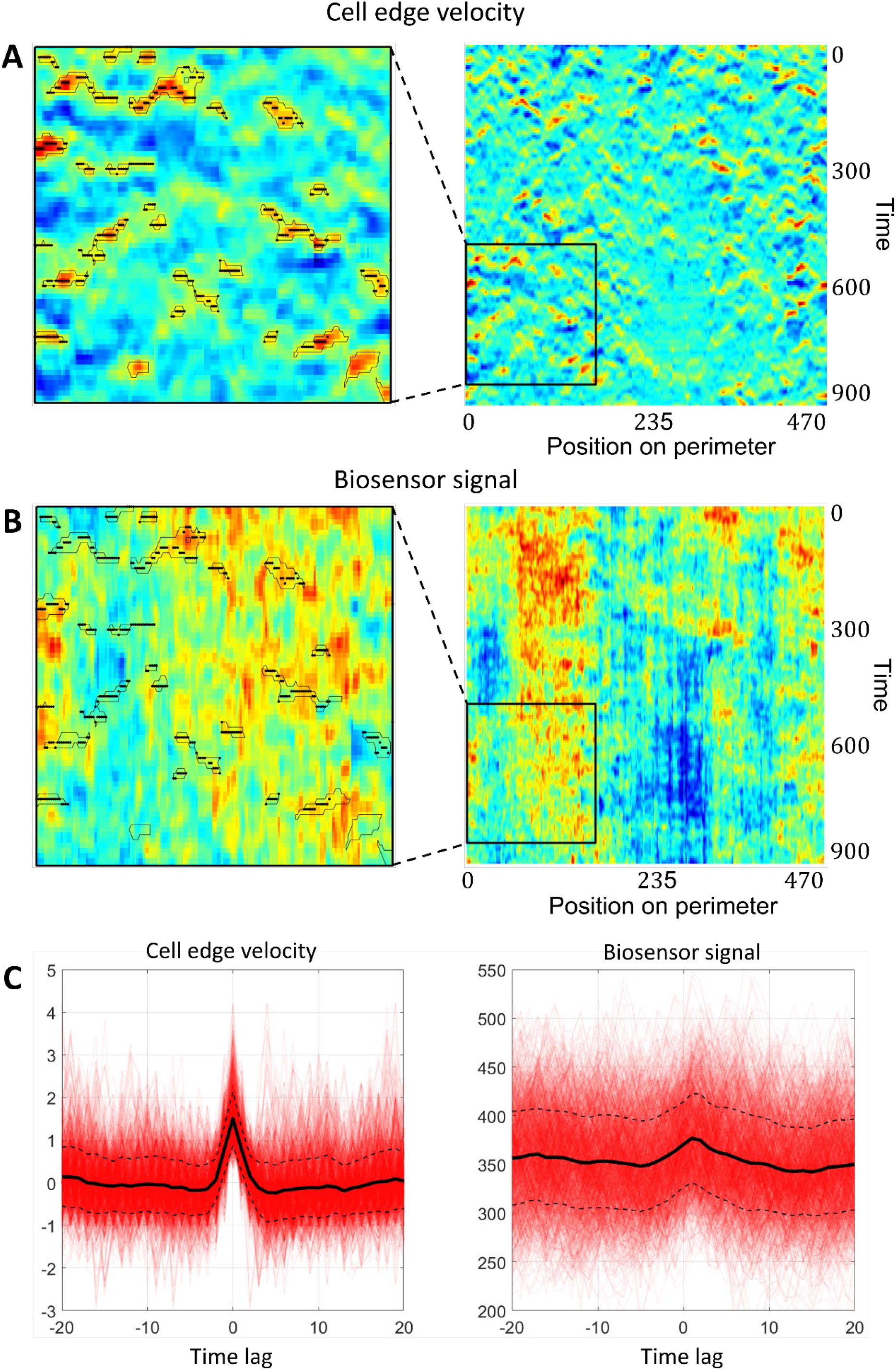
Simultaneous analysis of cell edge velocity and biosensor signal in the neighborhood of local maxima of cell edge velocity. **A**. In the obtained velocity kymographs we defined the areas of increased velocity values (black outlines) and found the points of local maximum within these regions (black dots). **B**. The points of local maximum were transferred from the velocity kymograph to the kymograph with biosensor signal kymograph. **C**. Both for velocity and for biosensor signal the values in the neighborhood of the identified points were extracted from kymographs and averaged to obtain the relative temporal comparison.

### The cell edge velocity is regulated by the GTPase’s rate of activation rather than its concentration value

Using the developed image analysis pipeline, we analyzed the dynamics of the cell membrane and GTPase activation in experimental data from breast cancer cells (MDA-MB-231). Both for Cdc42 and Rac1, the peaks of activity followed the peak of membrane velocity (**Fig. 5A-D, left two columns**). Given that actin polymerization and membrane protrusion are regulated downstream of Rac1 and Cdc42, such results may look counterintuitive. This effect was reported previously in several studies [17, 18, 21]. Yamao *et al*. suggested that the response function of a biosensor signal representing the regulation of membrane velocity has the properties of a differentiator circuit. However, the mechanism of such regulation is still not understood. To provide insight into this phenomenon, we first analyzed the dependence of membrane position (rather than velocity) on the biosensor signal and also the dependence of membrane velocity on the temporal derivative (i.e., the rate of change) of the biosensor signal. Membrane coordinate increased along the outline trajectories simultaneously with the biosensor signal (**Fig. 5E**). Membrane velocity decreased simultaneously with the temporal derivative of the biosensor signal (**Fig. 5F**). The time shift of Rac1 and Cdc42 peaks relative to the velocity peak was identical (each ∼5 seconds), which implies simultaneous activation of both GTPases (which is consistent with the previous study [17]). Next, we sought a simplest model that reproduces the temporal properties of Rac1 and Cdc42 activation in our 2D simulations of cell morphodynamics. We applied the same image analysis pipeline to the output (time-lapse images) of the considered models. For the first model setup (see **Methods** Eq. 11-14), we assumed the actin factor in the model is regulated by the concentration values of the active form of GTPases (**Fig. 5**, third column). In this case, the peak of velocity was closely aligned with the peak of GTPase concentration, the increase of the membrane position along trajectories followed the GTPase concentration peak, and the increase of the temporal derivative of the concentration preceded the membrane velocity increase (**Fig. 5D-F**, red arrows). Thus, none of the three quantitative characteristics agree with the experimental data. As an alternative model setup, we made the regulation of the actin factor dependent on the kinetic rate of GTPase activation (**Fig. 5**, rightmost column). In contrast to the first setup, we now reproduced the temporal shift of GTPase activity relative to the velocity peak, as well as the simultaneous increase of membrane position with the GTPase concentration and the simultaneous decrease of the membrane velocity with the decrease of the temporal derivative of the GTPase concentration (**Fig. 5D-F**, green arrows). These results suggest that membrane velocity is regulated by the kinetic rate of GTPase activation but not by the concentration of active GTPase. We interpret this finding in the following way: to power membrane protrusion during cell ruffling, it is not sufficient to maintain a certain level of GTPase activity. Instead, the membrane continues to protrude if the GTPase activity continues to increase. Once a max level of GTPase activity is reached, the protrusion stalls and the retraction cycle is initiated.

**Figure 5.**
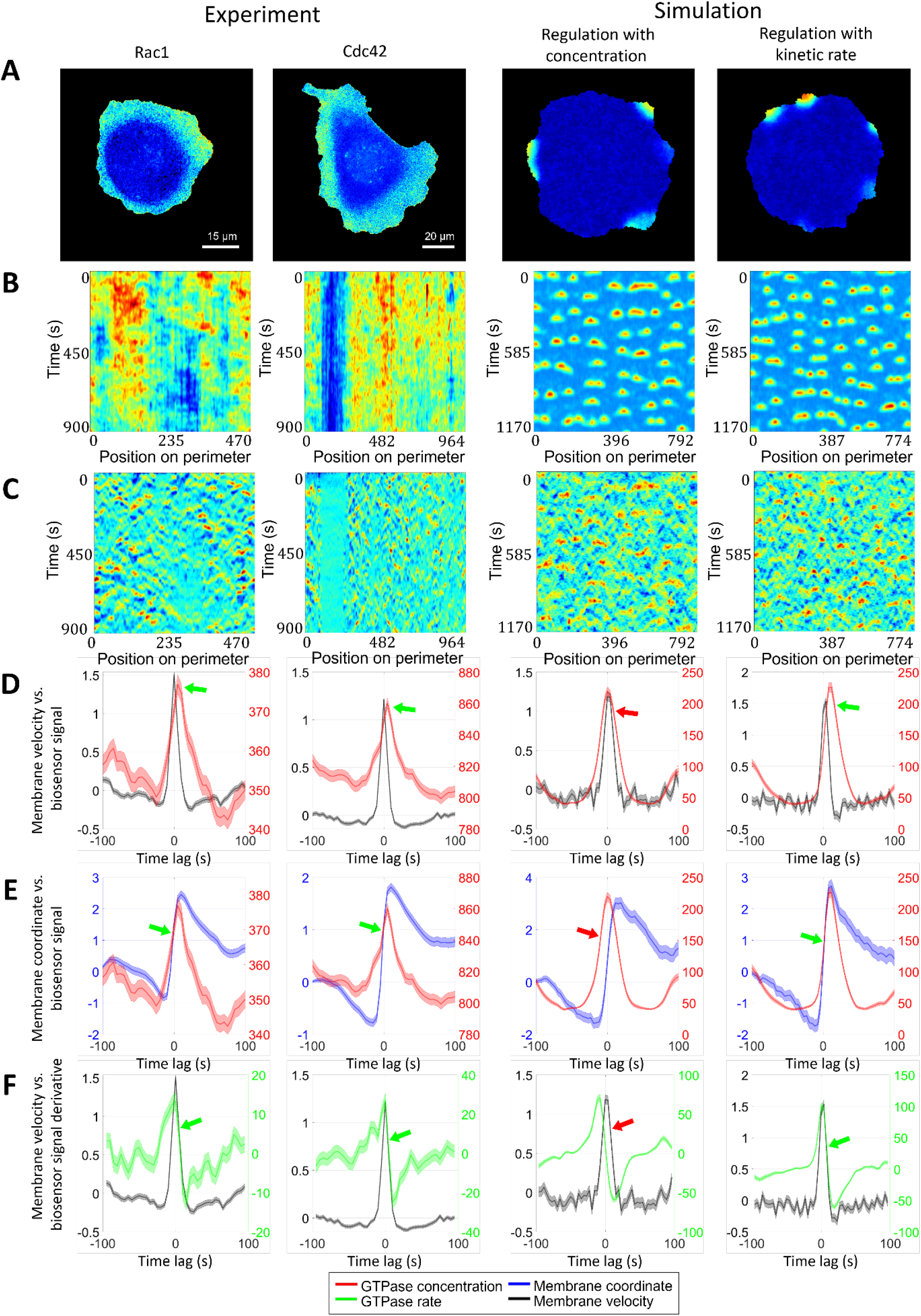
Regulation of GTPase activation by GTPase activation rate. We performed the same analysis for Rac1 and Cdc42 data in breast cancer cell line and in two simulation setups, where actin factor in the model is regulated by the absolute value of GTPase concentration and by the kinetic rate of GTPase activation. **A**. Snapshots of GTPase activity. **B**. Kymographs of GTPase activity. **C**. Kymographs of cell edge velocity. **D**. Comparison of cell edge velocity and biosensor signal (GTPase concentration). **E**. Comparison of cell edge coordinate and biosensor signal (GTPase concentration). **F**. Comparison of cell edge velocity and temporal derivative of biosensor signal (GTPase concentration). Features that are inconsistent with the experiment are shown with red arrows. Green arrows indicate agreement with the experiment.

### Cell-type specific relationship between peaks of Rac1 and Cdc42 activity can be reproduced with a unified model operating in different dynamic regimes

So far, we have focused on the analysis and modeling of GTPases activity during the ruffling of breast cancer cells (**Fig. 6A**). In this cell type, the peaks of Rac1 and Cdc42 activity occur simultaneously (i.e., with zero time delay). However, in mouse embryonic fibroblast (MEF) cells, FRET biosensor data [47] showed a small but distinctive time lag between the activation of the two GTPases. Peaks of Cdc42 activity precede peaks of Rac1 activity by 5 seconds on average (**Fig. 6B**). To understand the temporal regulation of GTPase activity, we considered several minimal models and explored the possibility to reproduce different time delays between Rac1 and Cdc42. We assumed that the time delay could be controlled by modulating the parameters involved in the feedback loops between the signaling components and by the sensitivity of Rac1 and Cdc42 to the upstream effector.

**Figure 6.**
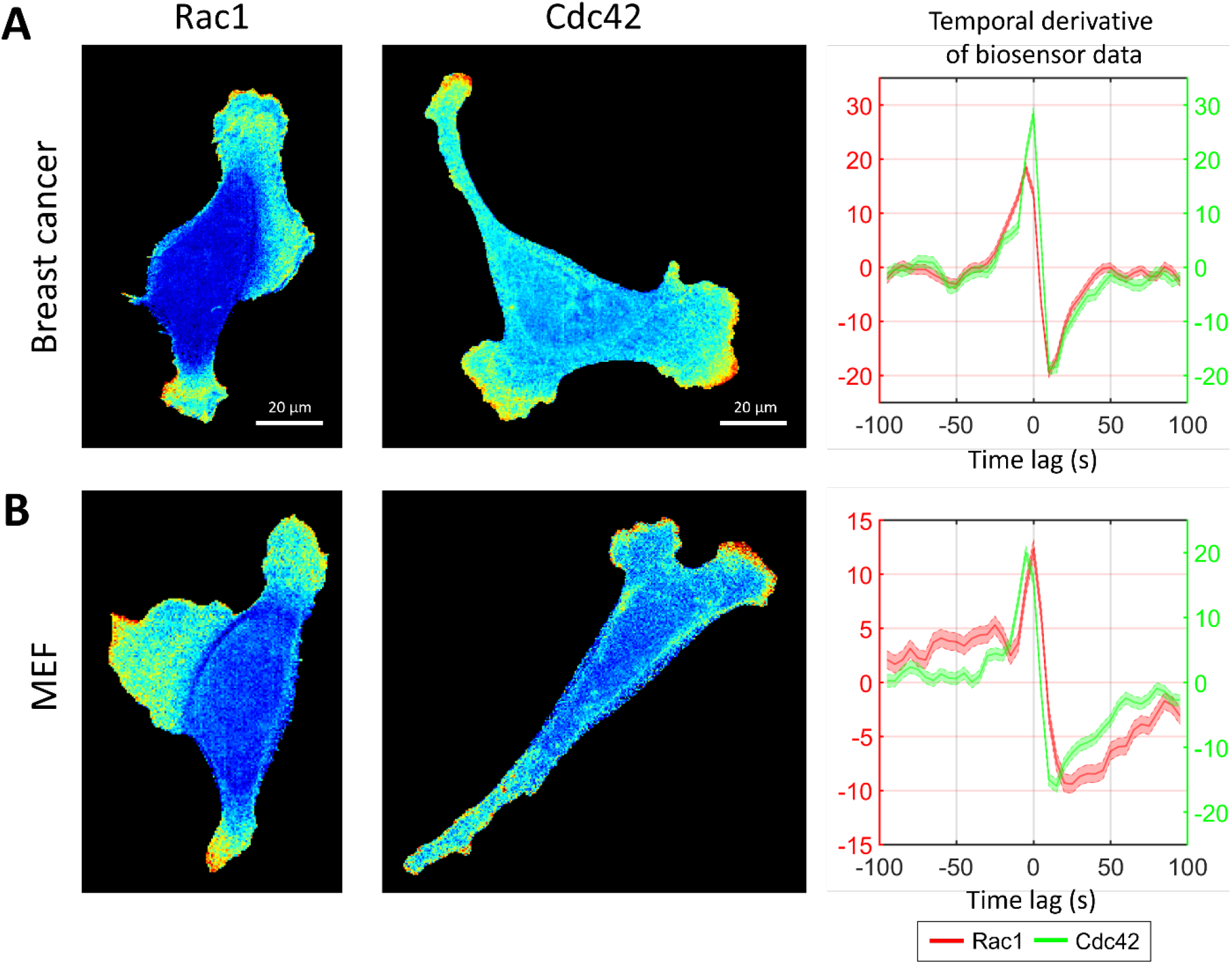
Simultaneous and delayed activation of Rac1 and Cdc42. **A**. Breast cancer cell line shows coherent activation of Rac1 and Cdc42. **B**. A delay between Cdc42 and Rac1 is observed in MEF cell line.

We considered models with a ‘crosstalk’ between Rac1 and Cdc42, as was previously reported in several studies [7, 12, 56]. These experiments indicate that Rac1 activates in response to the induced activation of Cdc42. Thus, in our model, we assumed positive regulation of Rac1 by Cdc42. However, the opposite regulation (from Rac1 to Cdc42) cannot be excluded because such bidirectional crosstalk is possible through the interaction of Rac1 and Cdc42 with their common GEFs [7, 57].

We first considered a model where Rac1 activity is induced by Cdc42 (**Fig. 7A**, Methods Eq. 15-20). Here the Rac1 component is represented with a bistable MCRD motif. The activation of Rac1 requires a transition from one stable state to the other stable state, which takes place when the active Cdc42 level reaches a certain threshold (**Supplemental Figure 3**). In this case, Rac1 activation is always delayed relative to the peak of Cdc42 activity. We conclude that such a model is consistent with MEF data but cannot reproduce the simultaneous activation of GTPases in breast cancer cells.

**Figure 7.**
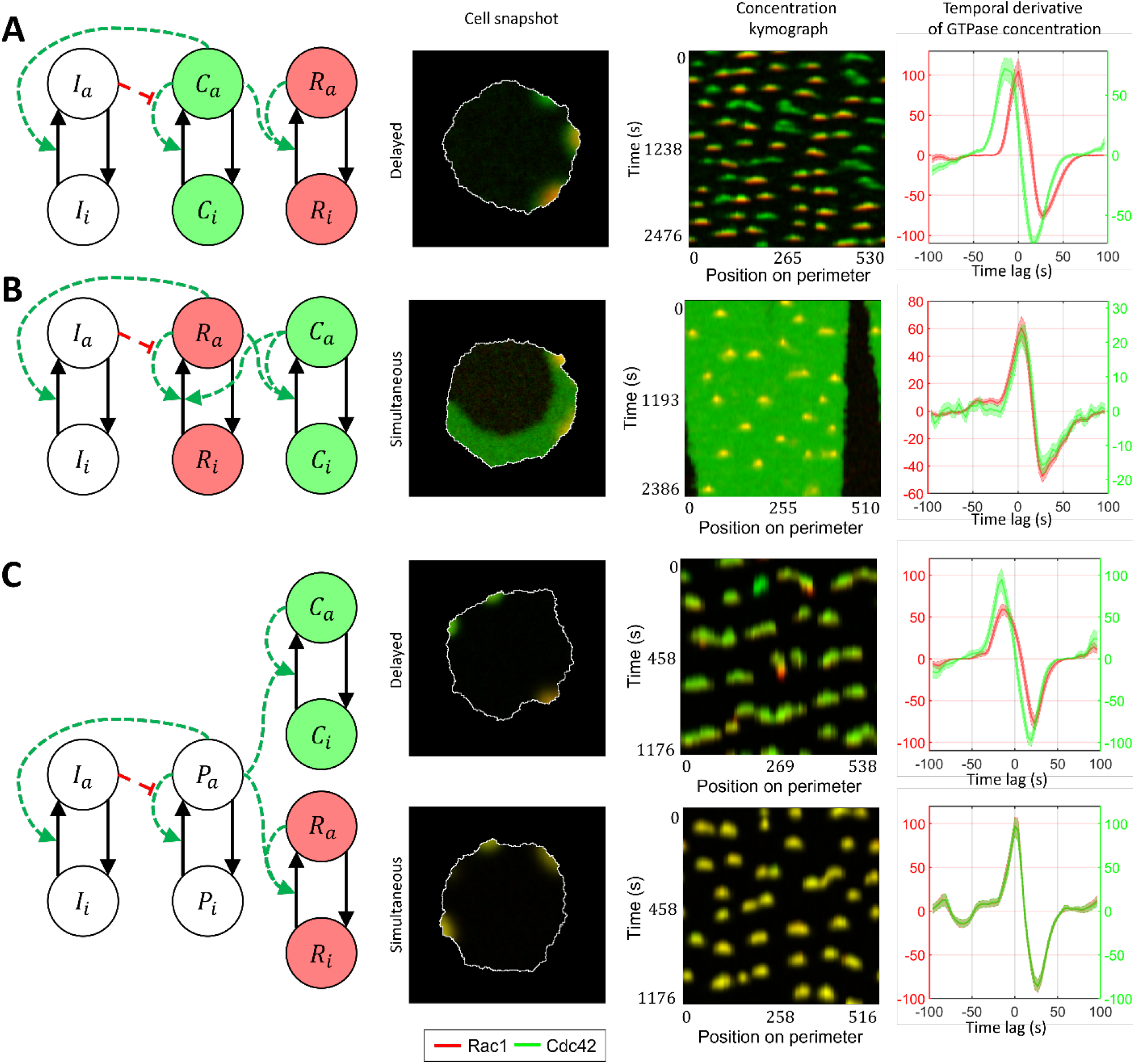
Signaling motifs including both Rac1 and Cdc42. **A**. Model of Rac1 activation induced by Cdc42 (leads to delayed activation of Rac1). **B**. Bidirectionally coupled model of Cdc42 and Rac1 activation (leads to simultaneous activation of GTPases). **C**. Model of Cdc42 and Rac1 induced by the upstream effector (can lead both to simultaneous and delayed activation).

As a next level of complexity, we consider a model where both GTPases are represented with bistable motifs bidirectionally coupled to each other (**Fig. 7B, Supplemental Video 1**). It was previously reported that Cdc42 has feedback on Rac1 [3, 12] and defines cell polarization [58, 59], while Rac1 drives protrusion and actin polymerization [33]. Thus, we sought to develop a model that captures these roles of Rac1 and Cdc42. Specifically, we expect Cdc42 in our model to generate polarization patch(es) at the cell edge. This way, through the feedback on Rac1, Cdc42 defines the parts of the cell periphery where the activity of Rac1 can drive the protrusion/retraction cycles. Because of the opposite feedback on Cdc42, Rac1, in turn, affects the variation of Cdc42 activity in protrusions. In this case, the increased activation of Cdc42 does not require its transition to a new stable state, and Cdc42 concentration varies near the level of a high-activity stable state. Such increased activity of Cdc42 above the high activity stable state is possible only if the crosstalk between Cdc42 and Rac1 is relatively weak. Mutual activation of two GTPases works as positive feedback, which can switch the system to a state where both Rac1 and Cdc42 can only be in the active state (**Supplemental Figure 4**). Such a state represents a polarized state of the cell. Although this model accurately reproduces the typical polarized ruffling dynamics as observed in breast cancer cells (MDA-MB-231), it only reproduces simultaneous but not the delayed activation of Rac1 and Cdc42. Therefore, the model in this form fits breast cancer cell data but not MEF cell data.

Next, as an alternative to the cell ruffling model that relies on feedback between Cdc42 to Rac1, we considered a model where instead of feedback regulation between Cdc42 and Rac1, both GTPases are activated in response to an upstream stimulus that drives their dynamics (**Fig. 7C, Supplemental Video 2**). Such upstream signaling motif could work through the PI3K pathway, which was reported as a regulator of cell ruffling [38] and can activate Rac1 and Cdc42 through the interactions of their GEFs with phosphoinositides [13]. This model allowed us to obtain both simultaneous activations of GTPases (when the response of the positive feedback in GTPase activation to the upstream stimulus is the same for Rac1 and Cdc42) and the delayed activation of Rac1 (when the threshold of activation for Rac1 was higher than for Cdc42). In the latter case, the difference in the activation thresholds is modulated by the inflection and max values of the Hill function representing the positive feedback (**Fig. 7C**).

Finally, we investigated the role of the crosstalk between Cdc42 and Rac1 in the model described in Fig.7C (with the upstream regulator). We found out that even if the responses of Rac1 and Cdc42 to the upstream effector are different and there is a delay in Rac1 and Cdc42 activation (Cdc42 precedes Rac1), the feedback from Cdc42 to Rac1 can compensate for this delay creating the simultaneous activation of the two GTPases (**Fig. 8A, Supplemental Video 3**). We quantified this effect in the 1D model for various values of *μ*_7_ parameter (**Fig. 8B**). As the strength of the feedback from Cdc42 to Rac1 increases, the delay between Cdc42 and Rac1 becomes negligible. Thus, we conclude that the unified model with the upstream effector motif and the feedback from Cdc42 to Rac1 can explain both the simultaneous dynamics of Cdc42 and Rac1 in the breast cancer cell line (MDA-MB-231) and the delayed activation in the MEF cell line.

**Figure 8.**
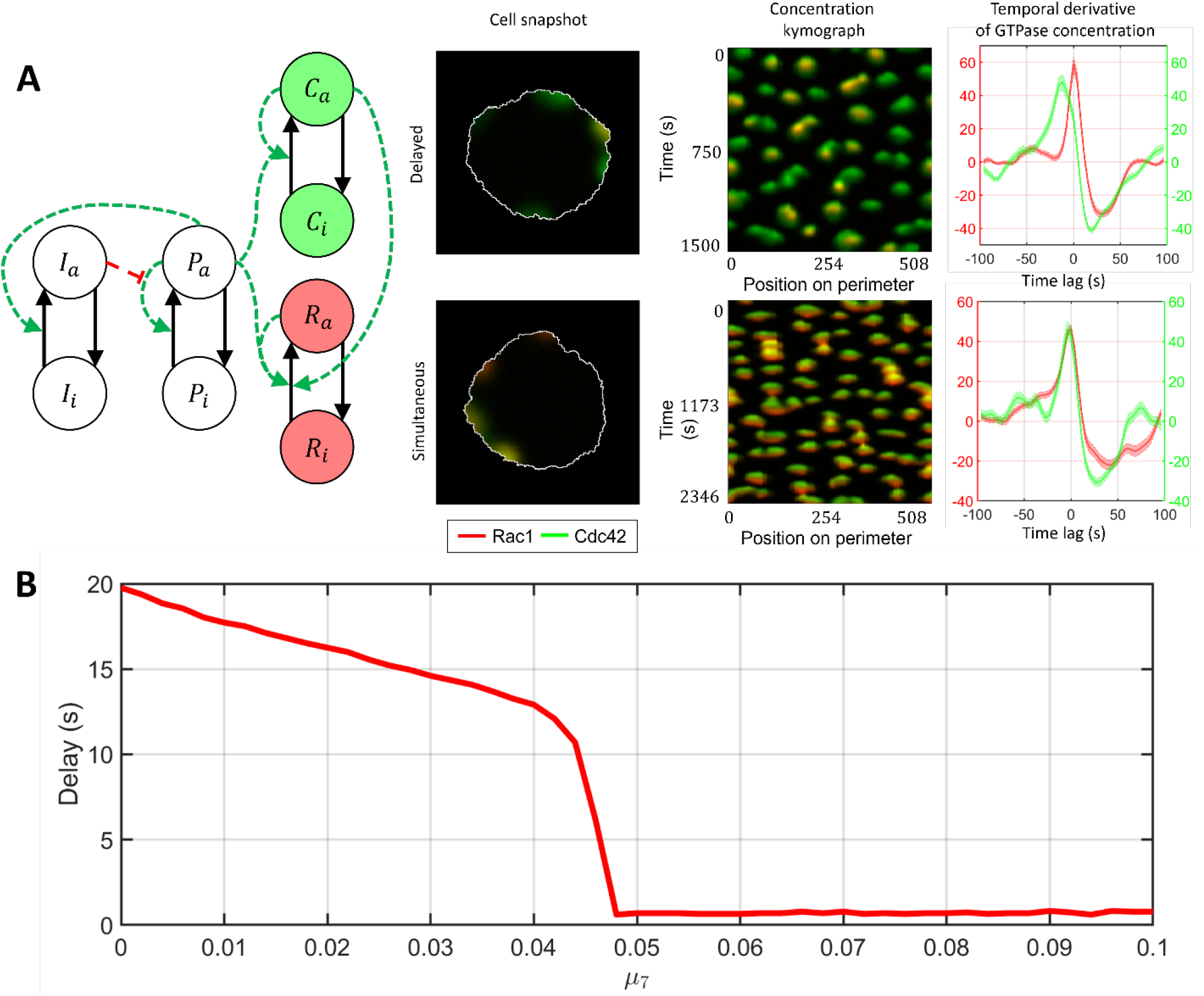
A unified model that accounts for upstream regulator of Cdc42 and Rac1 and for feedback between Cdc42 and Rac1. **A**. Signaling motif and two cases of dynamics: delayed and simultaneous activation of Rac1 and Cdc42. **B**. Modulation of feedback between Cdc42 and Rac1 can compensate the time delay between two GTPases.

## Discussion

Cell morphodynamics is a complex, multiscale process that involves regulation at different stages, from the biochemical regulation of protein activity to the biomechanical regulation of membrane protrusion through the cytoskeleton assembly. Understanding this process requires an integrative approach that connects multiple levels of regulation and represents them in a holistic manner. Computational modeling can be particularly useful for exploring the underlying processes. In the form of an in-silico experiment, computational modeling can be used to test conceptual biological hypotheses about regulatory mechanisms and to provide experimentally testable predictions.

In this study, we built our model to capture spatial and temporal GTPase activity and couple it to the protrusion/retraction cycles of cell edge motion along the whole cell periphery. We represented protein activity in the form of reaction-diffusion equations, which allowed us to apply the principles of pattern formation in biological morphogenesis to cellular-level dynamics and investigate the spatiotemporal activity of GTPases. The output of our model has the same format as the experimental imaging data, which enables a direct quantitative comparison of our simulation results and the data.

To perform the coordinated analysis of GTPase activity and cell edge motion, we developed an automated image analysis pipeline that computes a set of trajectories formed by the edge motion and tracks both GTPase activity and the edge velocity along these trajectories. We used this pipeline to match GTPase dynamics from experimental FRET biosensor data and from our simulation models. It allowed us to achieve close quantitative agreement of the modeled and observed dynamics of Cdc42 and Rac1 activity during cell ruffling in both the breast cancer cell (MDA-MB-231) and the MEF cell lines.

We tested two hypotheses about the regulation of protrusive activity. In one case, we assumed that the presence of active GTPase (e.g., Rac1) is sufficient to maintain the protrusion process, i.e., the cell edge responds to the concentration value of active GTPase. In the second case, we assumed that the increase of GTPase activity is needed to maintain the protrusion process, i.e., the cell edge responds to the kinetic rate of GTPase activation. We found out that the first hypothesis did not agree with the experimental data, but the second one showed close agreement with all metrics. One possible interpretation of this result is that maintaining a high level of GTPase activity may not be enough to drive the continuous protrusion and may lead to stalling, while the change in GTPase activity signals the cell to protrude or retract. Another possible interpretation is that our results reflect the fact that GTPase activity is powered by GTP hydrolysis, and thus the protrusion cycle may be synchronous with the local influx of GTP. In any case, further investigations are needed to determine the specific mechanism of such rate-of-change regulation.

Another focus of our morphodynamic cell modeling effort was to investigate the coordination between Rac1 and Cdc42 activity during cell ruffling. In addition to the breast cancer cell line (MDA-MB-231), where both GTPases are activated simultaneously, we also applied our image analysis pipeline to the MEF cell line. Since the delay in the peaks of GTPase activity can be cell-type specific (and our data analysis supports that possibility), we explored different models to test if the activity delay can be modulated within a unified regulatory network. We investigated different network motifs by feedback between Rac1 and Cdc42 and including an upstream effector acting on both Rac1 and Cdc42. We found that in the presence of the upstream regulator (presumably working through activation of phospholipids via the PI3K pathway), both simultaneous and delayed activations of Rac1 and Cdc42 are possible. In this setup, the feedback from Cdc42 to Rac1 can synchronize the activation of the two GTPases. An additional argument for the importance of the feedback between Cdc42 to Rac1 is that such feedback allowed us to capture polarized ruffling, i.e., a dynamic regime in which only one or several distinct parts of the cell undergo persistent ruffling. Our results provide insights into the regulatory mechanisms of GTPases and their role in cell morphodynamics, which can improve understanding of the underlying biological processes and explain differences in Rac1 and Cdc42 dynamics in various biological contexts. The scientific community can also benefit from our image analysis and modeling platforms in future studies of the mechanisms of cell motility driven by spatial and temporal dynamics of the regulatory proteins.

## Data Availability

MATLAB code for modeling and data analysis that was used in this study is available in the following GitHub repositories:

https://github.com/tsygankov-lab/Coupled_RDE-Morphodynamics_Model

https://github.com/tsygankov-lab/Cell_Ruffling_Quantification

## Authors Contributions

D.T. conceived the project. D.T., S.H., and S.N. conceptualized the study. S.H. and D.T. developed computational and image analysis approaches used in this work. S.H. designed and implemented simulation models and performed the comparative analysis of experimental data and simulation output. K.K. performed parameter optimization for some of the signaling motifs. J.S. and K.K. performed the textural analysis of the effect of noise on activity patterns. S.H. wrote the initial manuscript draft and prepared all figures and videos. S.H., D.T., S.N., and J.S. edited later manuscript versions. D.T. oversaw all aspects of the study.

## Acknowledgments

The authors would like to acknowledge the Partnership for an Advanced Computing Environment (PACE) at Georgia Tech and Dr. Melissa Kemp for the provided computational resources. The authors also thank Dr. Klaus M. Hanh, Dr. Daniel J. Marston, and the UNC-Olympus Research Imaging Center for providing the live-cell biosensor data used in this study.

## Supplementary Material

1. Supplemental Information as a single PDF file.
2. Supplemental Videos 1-3

## Supplemental Video Captions

**Supplemental Video 1**. An example of a simulation output for the model of cell ruffling with a bidirectional coupling of Cdc42 and Rac1 activity. Cdc42 defines cell polarization (green color) and the location where it induces Rac1 activity. Rac1 (red color) forms transient activity bursts, activating Cdc42 above the level in the polarization patch and driving cell protrusion.

**Supplemental Video 2**. An example of a simulation output for the model of cell ruffling where the external regulator induces Cdc42 (green) and Rac1 (red) activities. The movie shows two cases: delayed and simultaneous activation of Rac1 and Cdc42 due to a difference in the parameters of the response to the external regulator.

**Supplemental Video 3**. An example of a simulation output for the model of cell ruffling where Cdc42 (green) and Rac1 (red) activities are induced by the external regulator (as in Supplemental Video 2) but with additional feedback from Cdc42 (green) to Rac1 (red). The movie shows two cases: delayed and simultaneous activation of Rac1 and Cdc42 due to a difference in the strength of the feedback from Cdc42 to Rac1.

