## Supplemental Information for "Spatiotemporal coordination of Rac1 and Cdc42 at the whole cell level during cell ruffling"

**This PDF file includes:**

Supplemental Figures 1-4

Supplemental Text

**Other supplementary materials for this manuscript include:**

Supplemental Videos 1-3.

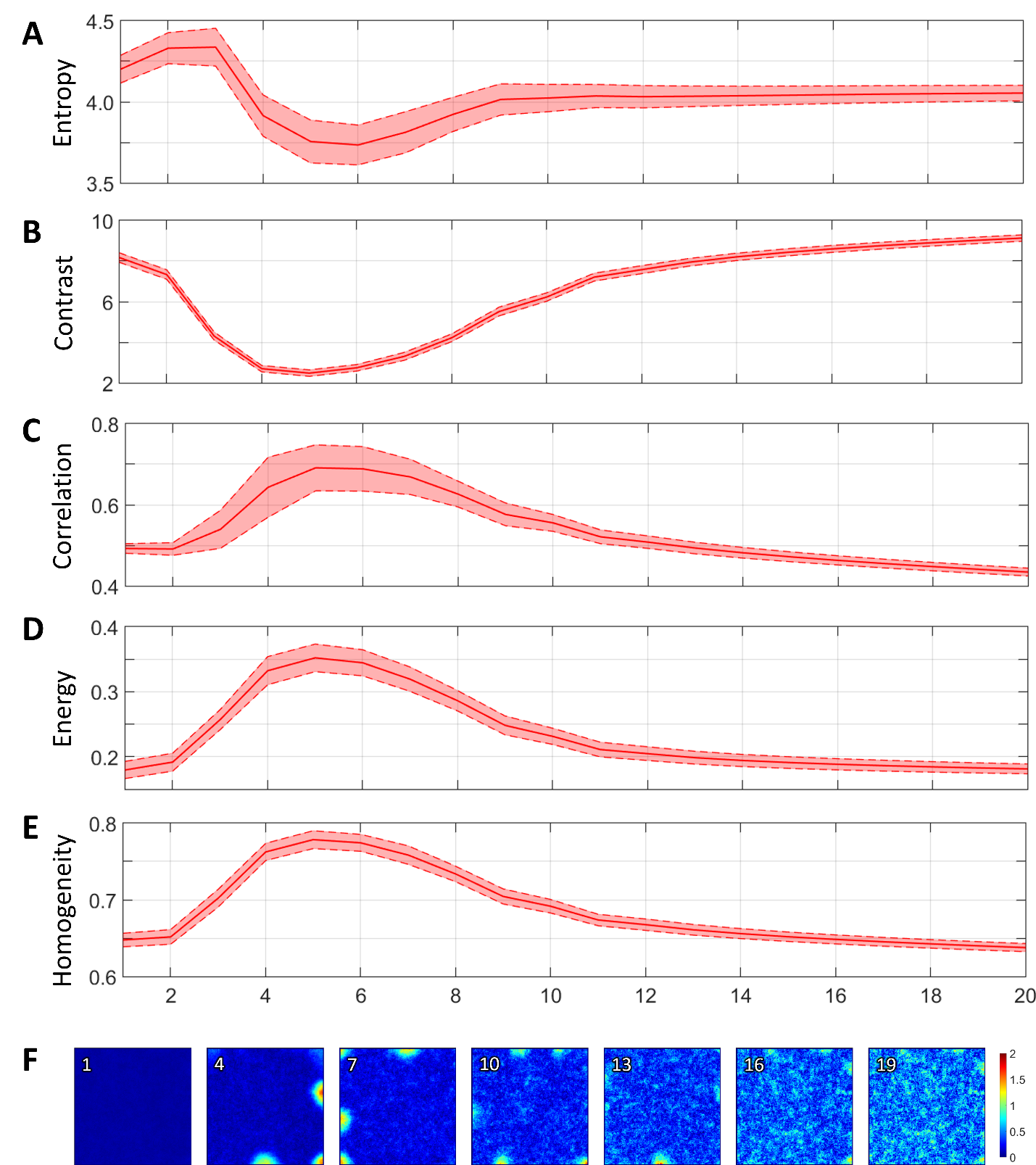

**Supplemental Figure 1**. Textural analysis of the model of GTPase activity in cell membrane ruffling for different values of noise. The values of kinetic parameters are the same as in **Fig. 2I**. For small values of noise no patterns were formed as diffusion fluxes did not allow amplification of GTPase at the perimeter. For increased values of noise patterns were formed along the perimeter. For higher values of noise (larger then 15) no patterns were observed because of too high value of noise. The formation of patterns can also be shown with the change of textural measures. **A**. Entropy. **B**. Contrast. **C**. Correlation. **D**. Energy. **E**. Homogeneity. **F**. Snapshots of simulation results for different values of noise.

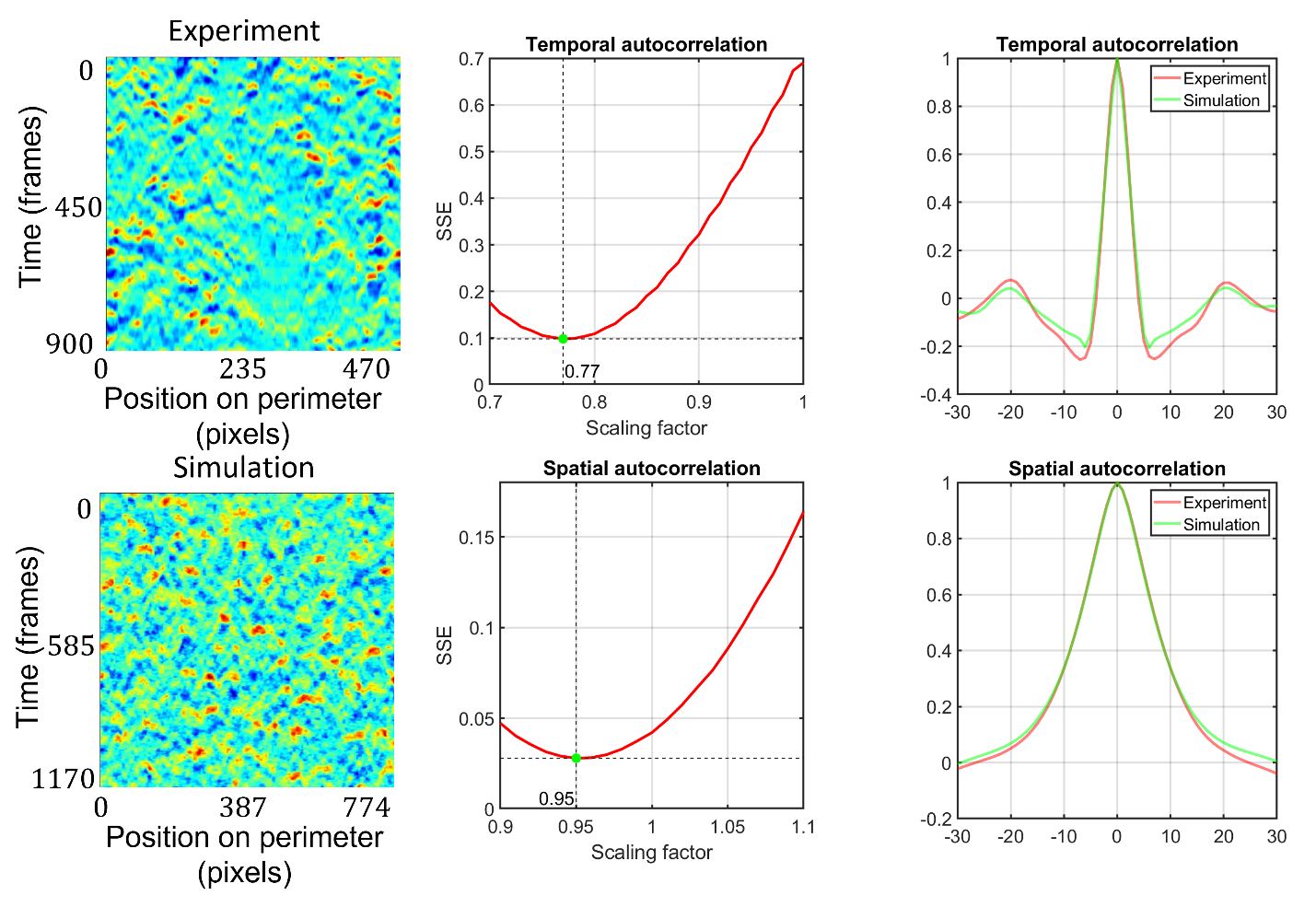

**Supplemental Figure 2**. Scaling of experimental and simulation data. Scaling was performed based on velocity kymographs using temporal and spatial autocorrelation plots. For a range of scaling coefficients for simulation we identified the factor with the minimal values of SSE (sum of squared errors).

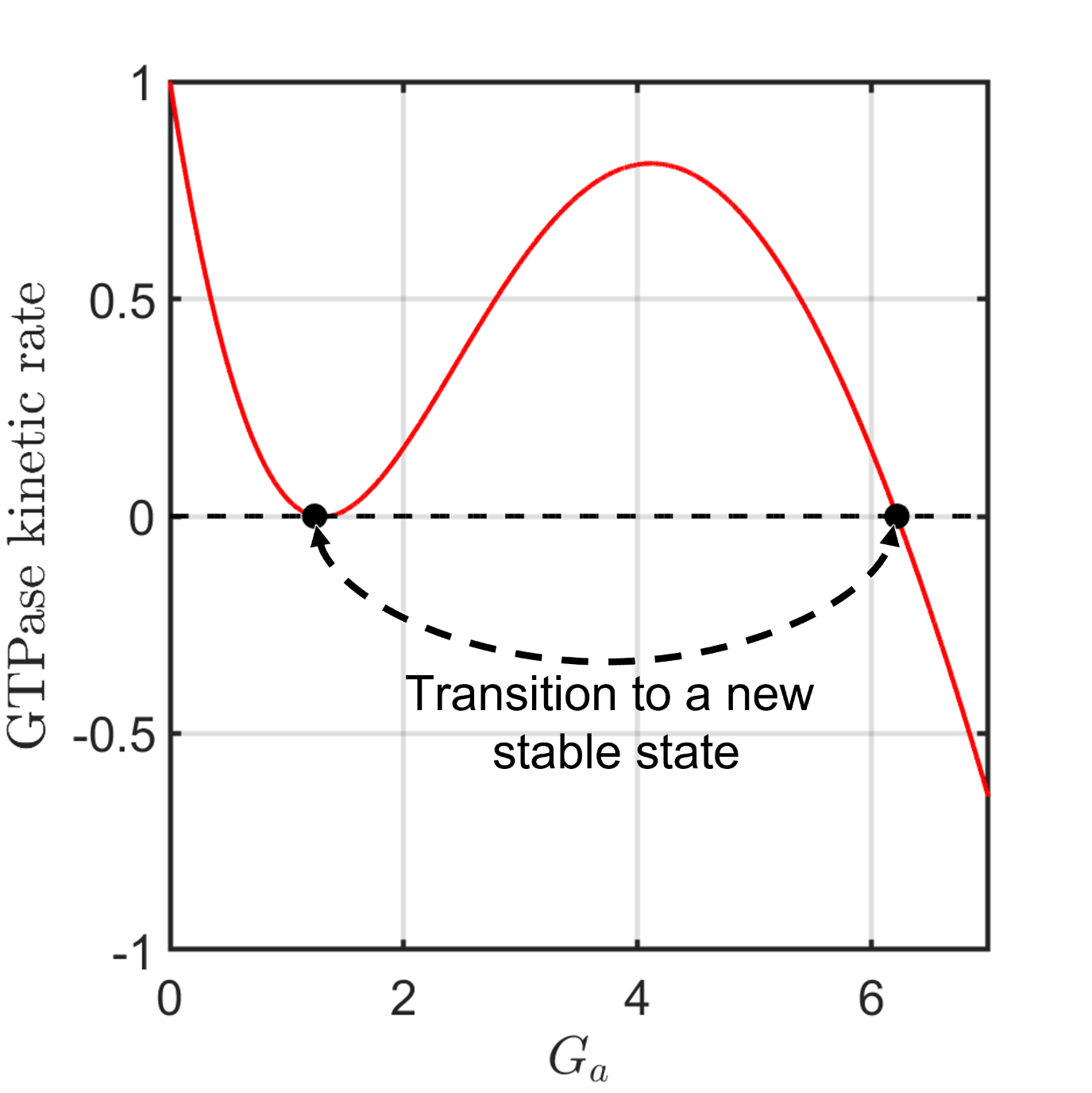

**Supplemental Figure 3**. Values of RHS (kinetic rate) function of GTPase activity during the transition to a new stable state. When the stimulus shifts the RHS so that homogeneous state is not stable (left), the transition to a new stable state is initiates, which requires time.

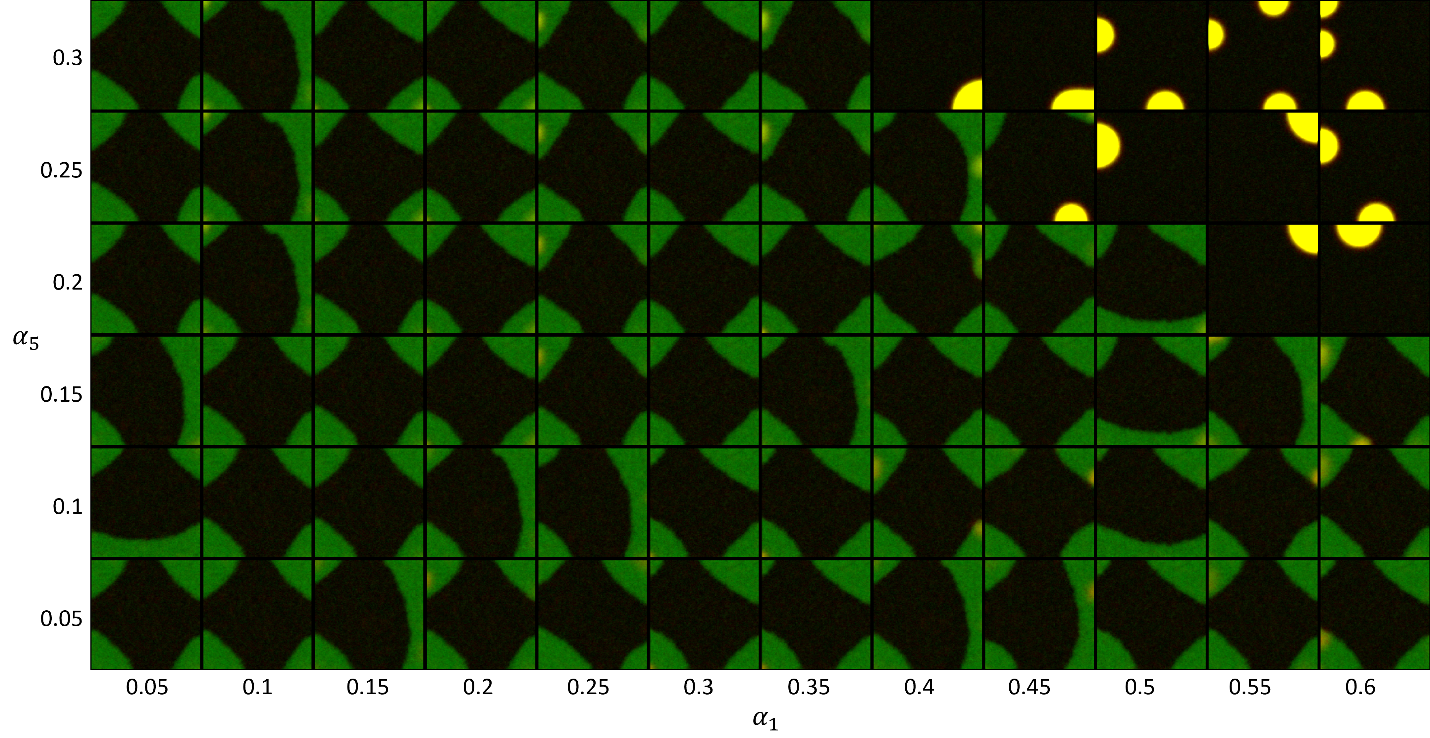

**Supplemental Figure 4**. Parameter scan for different values of feedbacks from Cdc42 to Rac1 ($\alpha_{1}$) and from Rac1 to Cdc42 ($\alpha_{5}$) in the model of bidirectionally coupled activity of Rac1 and Cdc42. For high enough values of feedbacks (upper right angle of the plot) the system switches from the dynamic regime of the formation of transient patches to the static regime, where activity of both GTPases are co-localized. Such state ca represent cell polarization.

**Supplemental Text**

To estimate the effect of GEF activation and deactivation rates on the GTPase activity, we considered a well-mixed system without diffusion. We first considered an extended mode of the two-component mass-conserved reaction-diffusion model, where the autocatalytic activation is represented explicitly: active GTPase positively regulates the rate of GEF activation, and activated GEF increases the rate of GTPase activation.

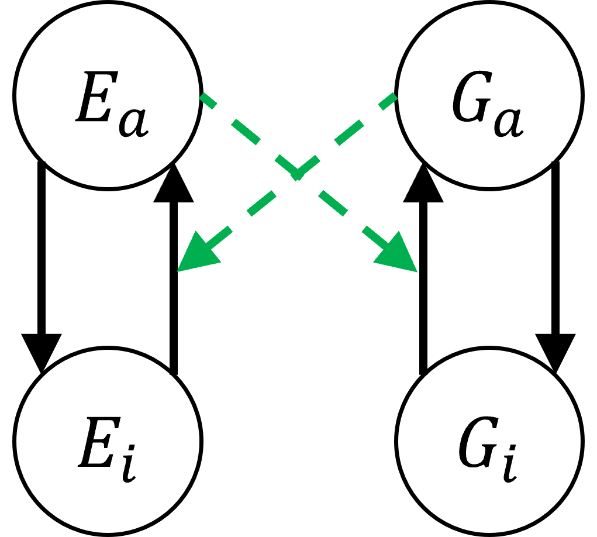
Inactive forms of the components ($E_{i}$, $G_{i}$) in the model were represented as a function of total concentration ($E_{T}, G_{T}$) and concentration of the active component ($E_{a}$, $G_{a}$):

$$E_{i}=E_{T}-E_{a}$$

$$G_{i}=G_{T}-G_{a}$$

$$\frac{\partial G_{a}}{\partial t}=\left( k_{1}+\gamma_{1}E_{a} \right)\left( G_{T}-G_{a} \right)-k_{2}G_{a}$$

$$\frac{\partial E_{a}}{\partial t}=\left( k_{3}+\gamma_{3}G_{a} \right)\left( E_{T}-E_{a} \right)-k_{4}E_{a}$$

For quasistatic assumption: $E_{a}\approx const$, so:

$$\left( k_{3}+\gamma_{3}G_{a} \right)\left( E_{T}-E_{a} \right)-k_{4}E_{a}=0$$

$$E_{a}=\frac{k_{3}+\gamma_{3}G_{a}}{k_{3}+k_{4}+\gamma_{3}G_{a}}E_{T}$$

If the effect of GTPase activation on GEF is significant in comparison to the basal activation of GEF, and the activation is fast, we can assume that $k_{3}\ll\gamma_{3}G_{a}$. In this case:

$$E_{a}\approx\frac{\gamma_{3}G_{a}}{k_{4}+\gamma_{3}G_{a}}E_{T}=\frac{G_{a}}{\frac{k_{4}}{\gamma_{3}}+G_{a}}E_{T}$$

If we use the notation: $\hat{\gamma}_{1}=\gamma_{3}E_{T}$ and $\hat{K}_{1}=\frac{k_{4}}{\gamma_{3}}$, we get:

$$\frac{\partial G_{a}}{\partial t}=\left( k_{1}+\frac{\hat{\gamma}_{1}G_{a}}{\hat{K}_{1}+G_{a}} \right)\left( G_{T}-G_{a} \right)-k_{2}G_{a}$$

If the response of GEF activation on active GTPase is nonlinear:

$$\frac{\partial E_{a}}{\partial t}=\left( k_{3}+\gamma_{3}G_{a}^{n} \right)\left( E_{T}-E_{a} \right)-k_{4}E_{a}$$

The approximated dynamics of GTPase will have the following form:

$$\frac{\partial G_{a}}{\partial t}=\left( k_{1}+\frac{\hat{\gamma}_{1}G_{a}^{n}}{\hat{K}_{1}^{n}+G_{a}^{n}} \right)\left( G_{T}-G_{a} \right)-k_{2}G_{a}$$

So, we can say that in the approximated system, the regulation of the GEF’s deactivation rate ($k_{4}$) corresponds to the regulation of the threshold of the autocatalytic activation ($\hat{K}_{1}$), while the magnitude of the positive feedback ($\hat{\gamma}_{1}$) is regulated by the total GEF concentration ($E_{T}$).

In the next step, we considered an extended system, where active GTPase increases the rate of inhibitor activation and inhibitor decreases the rate of GEF deactivation.

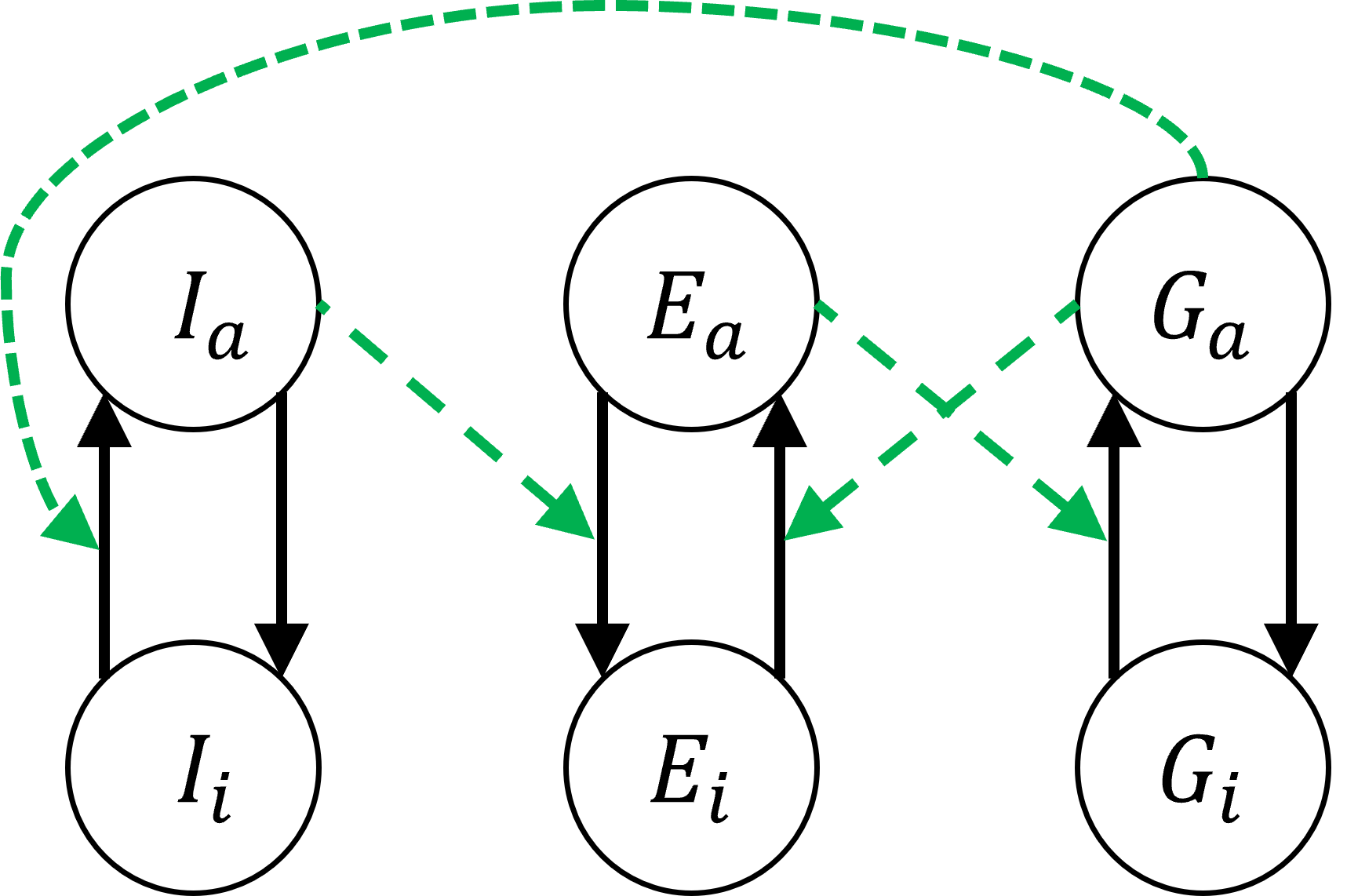
In this case, the well-mixed system is described by the following equations:

$$E_{i}=E_{T}-E_{a}$$

$$G_{i}=G_{T}-G_{a}$$

$$I_{i}=I_{T}-I_{a}$$

$$\frac{\partial G_{a}}{\partial t}=\left( k_{1}+\gamma_{1}E_{a} \right)\left( G_{T}-G_{a} \right)-k_{2}G_{a}$$

$$\frac{\partial E_{a}}{\partial t}=\left( k_{3}+\gamma_{3}G_{a} \right)\left( E_{T}-E_{a} \right)-(k_{4}+\gamma_{4}I_{a})E_{a}$$

$$\frac{\partial I_{a}}{\partial t}=\left( k_{5}+\gamma_{5}G_{a} \right)\left( I_{T}-I_{a} \right)-k_{6}I_{a}$$

For quasistatic condition $E_{a}\approx const$:

$$\left( k_{3}+\gamma_{3}G_{a} \right)\left( E_{T}-E_{a} \right)-(k_{4}+\gamma_{4}I_{a})E_{a}$$

$$E_{a}=\frac{k_{3}+\gamma_{3}G_{a}}{k_{3}+k_{4}+\gamma_{4}I_{a}+\gamma_{3}G_{a}}E_{T}$$

As previously, assuming that $k_{3}\ll\gamma_{3}G_{a}$:

$$E_{a}\approx\frac{\gamma_{3}G_{a}}{k_{4}+\gamma_{4}I_{a}+\gamma_{3}G_{a}}E_{T}=\approx\frac{G_{a}}{\frac{k_{4}}{\gamma_{3}}+\frac{\gamma_{4}}{\gamma_{3}}I_{a}+G_{a}}E_{T}$$

The dynamics of active GTPase is described in this case with the equation:

$$\frac{\partial G_{a}}{\partial t}=\left( k_{1}+\frac{\hat{\gamma}_{1}G_{a}}{\hat{K}_{1}+\beta I_{a}+G_{a}} \right)\left( G_{T}-G_{a} \right)-k_{2}G_{a}$$

Where $\hat{\gamma}_{1}=\gamma_{3}E_{T}$, $\hat{K}_{1}=\frac{k_{4}}{\gamma_{3}}$, $\beta=\frac{\gamma_{4}}{\gamma_{3}}$.
